# Telomere length sensitive regulation of Interleukin Receptor 1 type 1 (IL1R1) by the shelterin protein TRF2 modulates immune signalling in the tumour microenvironment

**DOI:** 10.1101/2021.12.07.471419

**Authors:** Ananda Kishore Mukherjee, Subhajit Dutta, Ankita Singh, Shalu Sharma, Shuvra Shekhar Roy, Antara Sengupta, Megha Chatterjee, Soujanya Vinayagamurthy, Sulochana Bagri, Divya Khanna, Meenakshi Verma, Dristhi Soni, Anshul Budharaja, Sagar Kailasrao Bhisade, Vivekanand, Ahmad Perwez, Nija George, Mohammed Faruq, Ishaan Gupta, Radhakrishnan Sabarinathan, Shantanu Chowdhury

## Abstract

Telomeres are crucial for cancer progression. Immune signalling in the tumour microenvironment has been shown to be very important in cancer prognosis. However, the mechanisms by which telomeres might affect tumour immune response remain poorly understood. Here, we observed that interleukin-1 signalling is telomere-length dependent in cancer cells. Mechanistically, non-telomeric TRF2 (Telomeric Repeat binding Factor 2) binding at the IL-1-receptor type-1 (IL1R1) promoter was found to be affected by telomere length. Enhanced TRF2 binding at the *IL1R1* promoter in cells with short telomeres directly recruited the histone-acetyl-transferase (HAT) p300, and consequent H3K27 acetylation activated IL1R1. This altered NF-kappa B signalling and affected downstream cytokines like *IL6, IL8* and *TNF*. Further, *IL1R1* expression was telomere-sensitive in triple-negative breast cancer (TNBC) clinical samples. Infiltration of tumour-associated macrophages (TAM) was also sensitive to the length of tumour cell telomeres and highly correlated with *IL1R1* expression. The use of both IL1 Receptor antagonist (IL1RA) and *IL1R1* targeting ligands could abrogate M2 macrophage infiltration in TNBC tumour organoids. In summary, using TNBC cancer tissue (>90 patients), tumour-derived organoids, cancer cells and xenograft tumours with either long or short telomeres, we uncovered a heretofore undeciphered function of telomeres in modulating IL1 signalling and tumour immunity.

## Introduction

Telomeres are composed of DNA repeats (TTAGGG) and associated ‘shelterin’ proteins present at the end of each eukaryotic chromosome. Gradual shortening of telomeres and change in the telomeric structure with cell division leads to cellular senescence/death, which is crucial for tissue homeostasis (Jacobs, 2013; Karlseder, 2002; Shay and Wright, 2000). During oncogenic transformation, the telomere shortening mediated cellular senescence is bypassed-resulting in uncontrolled cell proliferation and cancer progression; underlining the impact of telomeres in cancer (Blackburn, 2000; Hanahan and Weinberg, 2000; Maciejowski and Lange, 2017; Paeschke et al., 2010; Shay, 2016).

Shortening of telomeres with ageing contributes to compromised immune response (Goronzy et al., 2006; Hodes et al., 2002; Pawelec et al., 2014; Sansoni et al., 2008; Weng, 2012). However, despite its relevance in cancer prognosis and intervention (Artandi and DePinho, 2009; Aviv et al., 2017; Chew et al., 2012; F Balkwill et al., 2001; Greider, 1998; Hanahan and Weinberg, 2011; Wang et al., 2017), how telomeres influence tumour immune response is poorly understood.

Major advances show how immune cells including tumour-infiltrating-lymphocytes (TIL), tumour-associated-macrophages (TAM), natural-killer and dendritic cells impact cancer progression (Binnewies et al., 2018; Gonzalez et al., 2018; Jochems and Schlom, 2011; Salgado and Gevaert, 2021). TAMs - typically M2 macrophage-like – predominantly promote immunosuppressive tumour microenvironment (TME) through secretion of anti-inflammatory cytokines (Burkholder et al., 2014; Cassetta et al., 2016; DeNardo and Ruffell, 2019; Gonzalez et al., 2018; Pittet et al., 2022; SD et al., 2020), implicating the role of TAMs in aggressive tumour growth (Hu et al., 2015; Pittet et al., 2022).

Poor prognosis in triple-negative breast cancers (TNBC) [negative for estrogen receptor (ER-), progesterone receptor (PR-), and HER2 receptor(HER2-)] has been frequently associated with TAMs(Wang et al., 2022; Z. Y. Yuan et al., 2014). Multiple reports show M2-like TAMs to be the most abundant monocytic immune cells in TNBC and implicate their role in invasiveness and aggressive tumour growth (Burkholder et al., 2014; Jeong et al., 2019; Mantovani and Locati, 2013; Medrek et al., 2012; Oner et al., 2020; Shun et al., 2005; Z.-Y. Yuan et al., 2014).

How the telomere length (TL) of cancer cells affects macrophage infiltration has not been studied. Seimiya. H and colleagues noted that telomere elongation led to differential expression of genes associated with innate immunity(Hirashima et al., 2013). A published work on colorectal cancer showed the association of telomere length of cancer cells to immunity-related genes(Lopez-Doriga et al., 2018). Another recent article showed that telomere maintenance by Telomerase (the key enzyme responsible for telomere maintenance) and ATL (alternative telomere lengthening ) related mechanisms led to alteration of markers of TAM like CD163 in glioblastomas(Hung et al., 2016). Taken together, these suggest the possibility of molecular links between TAM infiltration and telomeres.

TRF2 (Telomeric Repeat binding Factor 2) is a canonical member of the telomere capping complex called ‘shelterin’ with reported extra-telomeric functions(Martínez and Blasco, 2011; Vinayagamurthy et al., 2020). We have reported earlier that the telomeric protein TRF2 alters gene expression in a telomere length-dependent fashion (Mukherjee et al., 2018). TRF2, although primarily a telomere-binding protein, was found to associate with gene promoters throughout the genome independent of its telomere-specific functions (Mukherjee et al., 2019a; Simonet et al., 2011; Yang et al., 2011). Interestingly, TRF2 was sequestered at the telomeres and away from gene promoters in cells with relatively longer telomeres, and vice-versa in the case of cells with shorter telomeres. This led to TRF2-mediated transcriptional regulation that was dependent on telomere length (a scheme of this model shown in **Additional Supplementary Figure 1**) (Mukherjee et al., 2018; Vinayagamurthy et al., 2023, 2020).

We found that non-telomeric TRF2 transcriptionally activated Interleukin-1-receptor type-1 (IL1R1) in a telomere-dependent fashion: resulting NF-kappaB (p65)-phosphorylation led to the upregulation of downstream cytokines. Telomere-sensitive IL1 signalling in TNBC was found to be correlated with enhanced TAM infiltration within the TME in tumours with relatively short telomeres. Together, these implicate telomere-sensitive immune-microenvironment in tumours.

## Results

### Non-telomeric TRF2 binding at the IL1 receptor type 1 (*IL1R1*) promoter is telomere length dependent

#### Cytokine and their receptor related genes are telomere-sensitive in cancer cells

We performed an RNA-seq experiment using our previously reported fibrosarcoma model of telomere elongation(Mukherjee et al., 2018). The transcriptomic profile of HT1080 cells (short telomere cells) was compared with HT1080-LT (long telomere cells). The difference in telomere length between the cells was confirmed using flow cytometry (**Supplementary Figure 1A**). The HT1080-LT cells had constitutively higher expression of *TERT* and *TERC* as previously described (**Supplementary Figure 1B**). The telomere length of the cells was further confirmed using *in-situ* hybridization of telomeric probes (**Supplementary Figure 1C**). Upon analyzing the RNA-seq data, we observed that several cytokine-related genes like *IL1B*, *IL1R1*, *TNF*, *IL15* and *IL37* had higher expression in HT1080 (short telomere) cells (**Supplementary Figure 1D**). We looked at the enriched pathways (KEGG) in genes that were upregulated (2 fold or higher expression) and down-regulated (0.5 fold or lower expression) in HT1080 cells (short telomere) compared to HT1080-LT (long telomere) cells (**Supplementary Figure 1E-F; Supplementary File 1**). It was observed that ‘Cytokine-cytokine receptor interaction’ was one of the top 5 most enriched pathways for the upregulated genes (**Supplementary Figure 1E**). While telomere elongation affected multiple pathways, we were most interested in cytokine-related genes that were differentially expressed due to the now well-established focus on the telomere-immunity axis(Barthel et al., 2017; Goronzy et al., 2006; Hirashima et al., 2013; Weng, 2012; Ye et al., 2014). We revisited our published ChIP-seq data in HT1080 fibrosarcoma cells(Mukherjee et al., 2019a) and found multiple cytokine-related gene promoters where TRF2 had peaks within 1000 bp from the transcriptional start site (TSS) (**Supplementary File 2**). We validated these sites by ChIP-qPCR in HT1080 fibrosarcoma cells and found that the *IL1R1* promoter had the most prominent TRF2 enrichment among the sites (**Supplementary Figure 1G**). We further performed ChIP-qPCR spanning +200 to -1000 bp of the *IL1R1* TSS to confirm promoter TRF2 binding across multiple cell lines-HT1080 fibrosarcoma, MDAMB231 breast cancer, HEK293T immortalized kidney and MRC5 fibroblast cells (**Supplementary Figure 1H**).

#### Telomere-dependent IL1R1 regulation in ex-vivo fibrosarcoma cells

Next, we tested the telomere dependence of TRF2 binding based on TL-sensitive non-telomeric TRF2 binding observed earlier (Mukherjee et al., 2018). TRF2-ChIP in ex-vivo cultured HT1080-LT/HT1080 cells showed that TRF2 occupancy at the *IL1R1* promoter was significantly reduced in cells with long telomeres relative to short telomere cells (**Figure 1A**). *IL1R1* mRNA expression was significantly lower in HT1080-LT relative to HT1080 cells (**Figure 1B**) conforming to the RNA-seq data described earlier (**Supplementary Figure 1D**). We found that the IL1R1 protein was also lower in HT1080-LT cells (flow cytometry; **Figure 1C**; Western Blot in **Supplementary Figure 1I**). Consistent with this, expression of IL1R1 at the cell surface was significantly lower in HT1080-LT compared to HT1080 cells (**Figure 1D**).

**Figure 1:**
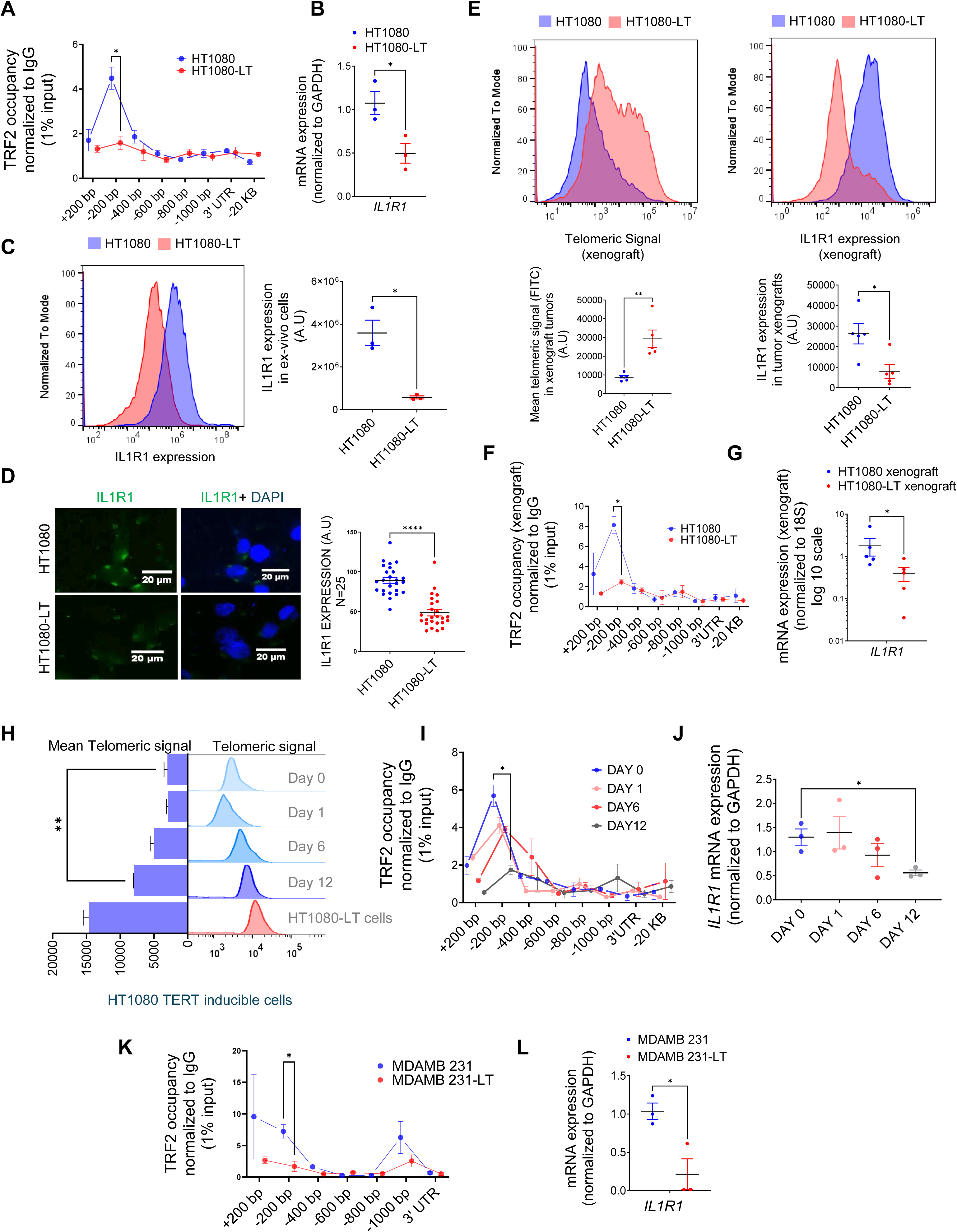
Expression of IL1R1 is regulated by telomere length-dependent enrichment of non-telomeric TRF2 on the gene promoter. A. Occupancy of TRF2 was checked by ChIP-qPCR on the *IL1R1* promoter spanning +200 to -1000 bp of TSS was reduced *ex-vivo* HT1080 cells with long telomeres (HL1080-LT) relative to ones with short telomeres (HT1080) cells (N=3 in each case) ;IL1R1-3’UTR or a region 20 kb upstream were used as negative controls for TRF2 binding. Statistical significance was calculated using unpaired T test with Welch’s correction (p values: * ≤ 0.05, ** ≤ 0.01, *** ≤ 0.001, **** ≤ 0.0001). B. mRNA expression of *IL1R1* in *ex-vivo* HT1080 or HT1080-LT cells; *GAPDH* was used for normalization [N=3]. Statistical significance was calculated using unpaired T test with Welch’s correction (p values: * ≤ 0.05, ** ≤ 0.01, *** ≤ 0.001, **** ≤ 0.0001). C. IL1R1 protein expression was checked by immuno-flow cytometry in HT1080 or HT1080-LT cells in three independent replicates; Mean IL1R1 expression has been plotted along the X-axis in log scale (right panel) [N=3]. Statistical significance was calculated using unpaired T test with Welch’s correction (p values: * ≤ 0.05, ** ≤ 0.01, *** ≤ 0.001, **** ≤ 0.0001). D. IL1R1 levels by immunofluorescence microscopy in HT1080 or HT1080-LT cells; cells were stained with DAPI for marking the cell nucleus; IL1R1 levels from 25 individual cells shown in graph (right panel). [N=25] Statistical significance was calculated using Mann-Whitney’s non-parametric test (p values: * ≤ 0.05, ** ≤ 0.01, *** ≤ 0.001, **** ≤ 0.0001). E. Telomere length of the xenograft tumors made in NOD SCID mice using HT1080 cells with long telomeres (HL1080-LT) and short telomeres (HT1080) was checked by flow-cytometry and was higher in HT1080-LT xenograft tumors. Following this, IL1R1 expression was checked by flow cytometry for these samples. Mean Telomeric signal and Mean IL1R1 expression has been plotted respectively ( panels below) [N=5] Statistical significance was calculated using Mann-Whitney’s non-parametric test (p values: * ≤ 0.05, ** ≤ 0.01, *** ≤ 0.001, **** ≤ 0.0001). F. Occupancy of TRF2 by ChIP-qPCR at the *IL1R1* promoter spanning +200 to -1000 bp of TSS was reduced in xenograft tumors made from HT1080 cells with long telomeres (HL1080-LT) relative to ones with short telomeres (HT1080) cells (N=3 in each case; IL1R1-3’UTR or a region 20 kb upstream were used as negative controls for TRF2 binding). [N=3] Statistical significance was calculated using unpaired T test with Welch’s correction (p values: * ≤ 0.05, ** ≤ 0.01, *** ≤ 0.001, **** ≤ 0.0001). G. mRNA expression of *IL1R1* in xenograft tumors (HT1080 or HT1080-LT cells); *18S* was used for normalization. [N=5] Statistical significance was calculated using Mann-Whitney’s non-parametric test (p values: * ≤ 0.05, ** ≤ 0.01, *** ≤ 0.001, **** ≤ 0.0001). H. Telomere length by flow-cytometry of telomeric signal in HT1080 cells (hTERT-inducible stable line) following 0, 1, 6, 12 or 20 days of hTERT induction; HT1080-LT shown as positive control for enhanced telomere length; mean telomeric signal (FITC) plotted bar graph (left). [N=3] Statistical significance was calculated using unpaired T test with Welch’s correction (p values: * ≤ 0.05, ** ≤ 0.01, *** ≤ 0.001, **** ≤ 0.0001). I. TRF2 occupancy ChIP-qPCR on the *IL1R1* promoter spanning +200 to -1000 bp of TSS in HT1080 cells following 0, 1, 6 or 12 days of hTERT induction in ex-vivo culture; primers and normalization as in Figure 2A; IL1R1-3’UTR or a region 20 kb upstream used as negative controls. [N=3] Statistical significance was calculated using unpaired T test with Welch’s correction (p values: * ≤ 0.05, ** ≤ 0.01, *** ≤ 0.001, **** ≤ 0.0001). J. *IL1R1* mRNA expression (normalized to *GAPDH*) in HT1080 cells following 0, 1, 6 or 12 days of hTERT induction in ex-vivo culture. [N=3] Statistical significance was calculated using unpaired T test with Welch’s correction (p values: * ≤ 0.05, ** ≤ 0.01, *** ≤ 0.001, **** ≤ 0.0001). K. TRF2 occupancy by ChIP-qPCR on the *IL1R1* promoter in MDAMB23 or MDAMD231-LT cells using primers and normalization described earlier. [N=3] Statistical significance was calculated using unpaired T test with Welch’s correction (p values: * ≤ 0.05, ** ≤ 0.01, *** ≤ 0.001, **** ≤ 0.0001). L. Expression for *IL1R1* in MDAMB231 cells or MDAMD231-LT cells; *GAPDH* was used for normalization. [N=3] Statistical significance was calculated using unpaired T test with Welch’s correction (p values: * ≤ 0.05, ** ≤ 0.01, *** ≤ 0.001, **** ≤ 0.0001). Error bars correspond to standard error of mean from independent experiments.

#### Telomere-dependent IL1R1 regulation in xenograft tumors

We generated xenograft tumours using HT1080 and HT1080-LT cells in NOD-SCID mice (see methods for details). We confirmed that there was higher expression of *TERT* and *TERC* in the HT1080-LT xenograft tumours (**Supplementary Figure 1J**). We found that the xenograft tumours from HT1080-LT cells had higher mean telomere length than those from HT1080 cells and had lower IL1R1 expression at the protein level (**Figure 1E**). It was observed that TRF2 occupancy (**Figure 1F**), as well as *IL1R1* mRNA (**Figure 1G**) expression, was lower in HT1080-LT-derived xenograft tumours.

#### TERT-inducible model of telomere elongation in fibrosarcoma cells

Telomere-dependent TRF2 binding at the *IL1R1* promoter was further checked using *hTERT*-inducible HT1080 cells that gradually increased telomeres by ∼50% over 12 days following hTERT induction (**Figure 1H**, **Supplementary Figure 1K**): gradual reduction of *IL1R1* promoter TRF2 binding (**Figure 1I**) and *IL1R1* expression (**Figure 1J**) with telomere elongation over the 12 days was notably consistent with observations made using HT1080/HT1080-LT cells (**Figure 1A-B**).

#### Telomere elongation model in MDAMB231 breast cancer cell line

An alternative telomere elongation model was made from TNBC (triple negative breast cancer) MDAMB-231 cells using G-rich-telomere-repeat (GTR) oligonucleotides as reported earlier (Mukherjee et al., 2018; Wright et al., 1996). In the resulting MDAMB-231-LT cells with ∼1.8-2-fold increase in TL (**Supplementary Figure 1L**), *IL1R1* promoter TRF2 occupancy and *IL1R1* expression were significantly lower (**Figure 1K-L**). The results collectively suggest that TRF2 occupancy at the *IL1R1* promoter and the expression of the gene is telomere length dependent.

### TRF2 transcriptionally activates *IL1R1*

We, next, investigated the role of TRF2 in the regulation of *IL1R1*.TRF2 over-expression (Flag-TRF2) showed an increase in *IL1R1* mRNA expression and protein, and TRF2 silencing reduced *IL1R1* in HT1080 cells (**Figure 2A**, and **Supplementary Figure 2A**) indicating that TRF2 activated *IL1R1* expression. Flag-tagged TRF2-full-length, or a truncated version of flag-tagged TRF2 protein without the necessary domains for DNA binding [N-terminal Basic-domain (TRF2-delB) or C terminal MYB-domain (TRF2-delM)], showed that both TRF2 occupancy and *IL1R1* promoter activity was compromised in case of the mutants; indicating TRF2 DNA binding as necessary for *IL1R1* activation (**Figure 2B-C**). We developed a doxycycline-inducible TRF2 model using HT1080 cells. Upon induction of TRF2, IL1R1 expression was also enhanced (**Figure 2D-E**). We checked and found that IL1R1 expression of the cell surface was also significantly higher in cells with higher TRF2 (**Figure 2F**). Further, TRF2-dependent activation of *IL1R1* was clear in MDAMB-231 cells (**Supplementary Figure 2B**), normal MRC5 fibroblasts and HEK293T cells (**Supplementary Figure 2C**). HT1080 cells with doxycycline-inducible-TRF2 were used to make tumour xenografts in NOD-SCID mice. TRF2-induced tumours had enhanced IL1R1 relative to the xenograft tumours where TRF2 was not induced (**Figure 2G**).

**Figure 2:**
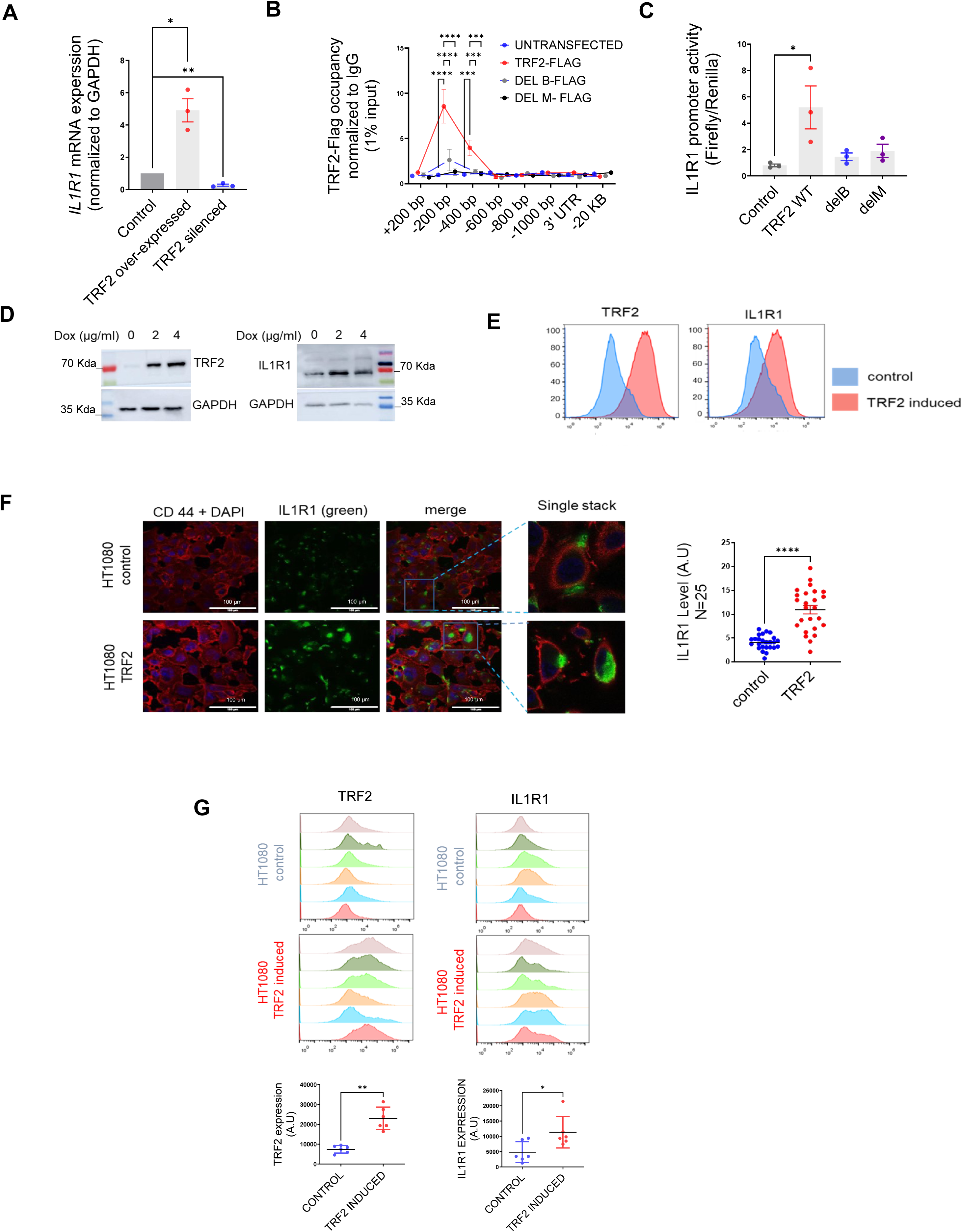
**TRF2 is a transcriptional activator of IL1R1** A. *IL1R1* expression by qRT PCR following expression of TRF2-flag or TRF2 silencing ( 48 hrs. of transient transfection) ; *GAPDH* was used for normalization. [N=3] Statistical significance was calculated using unpaired T test with Welch’s correction (p values: * ≤ 0.05, ** ≤ 0.01, *** ≤ 0.001, **** ≤ 0.0001). B-C. TRF2 ChIP-qPCR spanning +200 to -1000 bp of *IL1R1* TSS for TRF2 occupancy (B) and promoter activity (luciferase reporter including -1500 bp TSS; C) following expression of flag-tag TRF2 full-length, or the mutants TRF2-delB or TRF2-delM; anti-flag antibody used for ChIP; IL1R1-3’UTR or a region 20 kb upstream were used as negative controls. [N=3] Statistical significance was calculated using Tukey’s multiple comparisons test (p values: * ≤ 0.05, ** ≤ 0.01, *** ≤ 0.001, **** ≤ 0.0001). D. TRF2 and IL1R1 levels by western blots following TRF2 induction with 2 or 4 µg/ml doxycycline (Dox) in HT1080 cells (stably transformed for Dox-inducible TRF2); GAPDH as loading. E. TRF2 and IL1R1 expression in control or TRF2-induced HT1080 cells by flow-cytometry; TRF2/IL1R1 in x-axis in log scale for 20 thousand cells. F. IL1R1 levels in control or TRF2-induced HT1080 cells. CD44 used for cell-surface marker and nuclei were stained with DAPI; 25 cells in each condition were scored and plotted in the summary graph. [N=25] Statistical significance was calculated using Mann-Whitney’s non-parametric test (p values: * ≤ 0.05, ** ≤ 0.01, *** ≤ 0.001, **** ≤ 0.0001). G. TRF2 (left panel), IL1R1 (right panel) levels from xenograft tumors in NOD-SCID mice (control or doxycycline-induced TRF2 in HT1080 cells; N=6 mice in each group) by immuno-flow cytometry; mean fluorescence signal from individual tumors in control or TRF2-induced tumors plotted in adjacent graphs. [N=6] Statistical significance was calculated using Mann-Whitney’s non-parametric test (p values: * ≤ 0.05, ** ≤ 0.01, *** ≤ 0.001, **** ≤ 0.0001). Error bars correspond to standard error of mean from independent experiments.

### G-quadruplex DNA secondary structure necessary for TRF2 binding to the *IL1R1* promoter

We reported TRF2 binding to G-quadruplex (G4) DNA secondary structures within promoters throughout the genome (Mukherjee et al., 2019a); further, the role of G4s in epigenetic regulation was noted (Mukherjee et al., 2019b). Here, we found two G4s within the *IL1R1* promoter TRF2-binding site (**Figure 3A, Supplementary Figure 3A**). We confirmed that the DNA sequences formed G4 structures in solution and specific base substitutions disrupted the structure formation (**Figure 3B**); both G4 forming motif sequences showed the characteristic Circular Dichroism signatures for ‘parallel’ G-quadruplex structures which were largely abrogated when the sequences were mutated (**Figure 3B**). BG4 antibody (against G4s (Biffi et al., 2013)) ChIP was enriched ∼200 bp upstream on the *IL1R1* promoter in HT1080 cells(**Figure 3C**); TRF2-dependent *IL1R1* promoter activity was significantly compromised on introducing G4-deforming G˃T substitutions on both the G4 motifs (**Figure 3D**), while the effect of the mutation was more for G4 motif B. In the presence of the intracellular G4-binding ligand 360A (Granotier et al., 2005), the TRF2 occupancy in HT1080 cells was found to be significantly reduced at the *IL1R1* promoter consistent with results from other TRF2 binding sites (Mukherjee et al., 2019a) (**Supplementary Figure 3B**). Upon treatment of HT1080 cells with 360A, TRF2-induced IL1R1 promoter activity, and expression/protein were significantly reduced (**Figure 3E**). Using CRISPR, we artificially inserted an *IL1R1* luciferase-reporter-cassette (-1500 bp) at the CCR5 safe-harbour locus in HEK293T cells; TRF2-dependent promoter activity from the inserted *IL1R1* promoter-reporter and TRF2 binding was reduced when the G4 motif B was mutated supporting earlier results (**Figure 3F-G**). Furthermore, CRISPR-mediated G>T editing in the G4 motif (motif B) within the endogenous *IL1R1* promoter resulted in a reduction of TRF2 binding and *IL1R1* expression (**Supplementary Figure 3C-D**).

**Figure 3:**
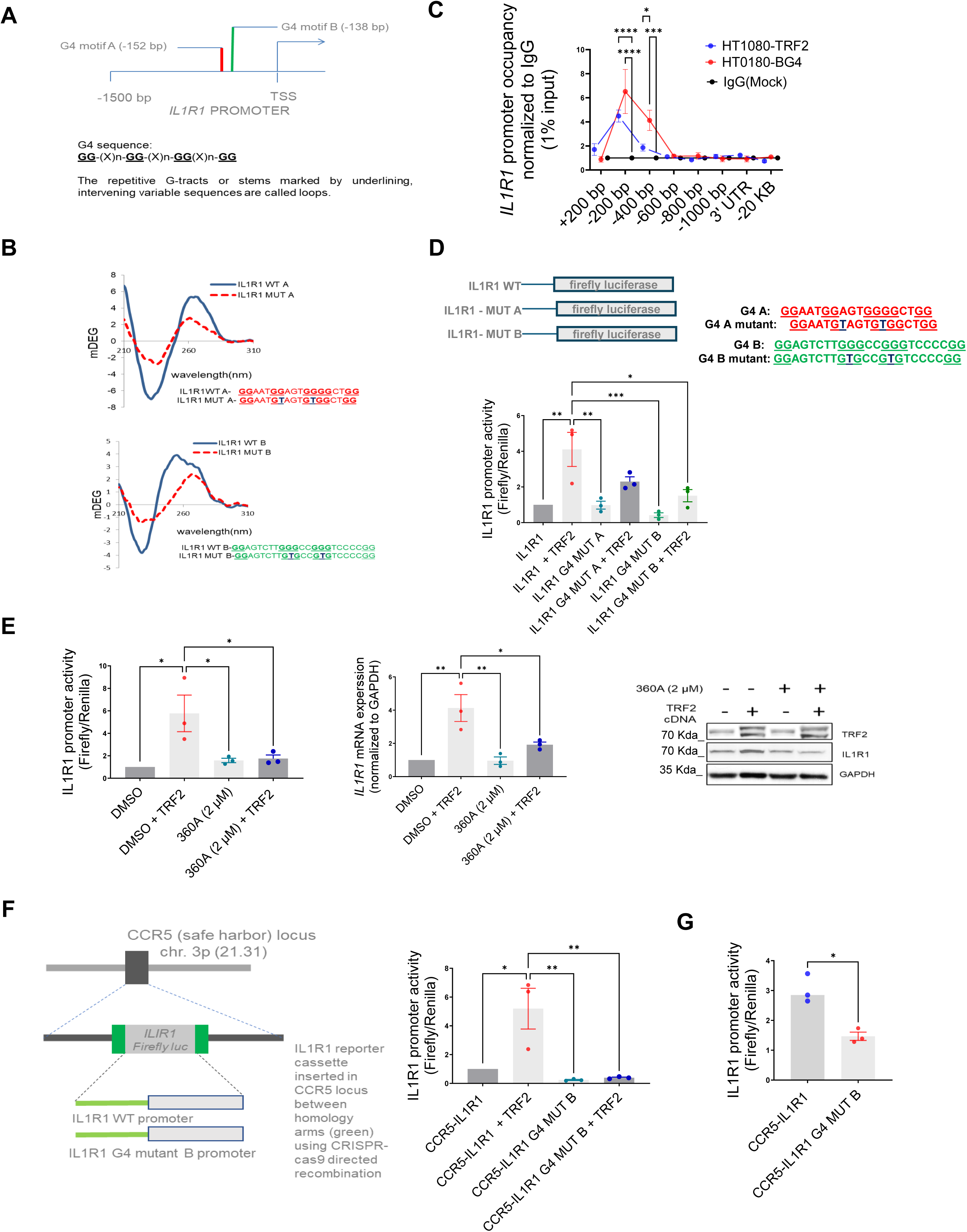
DNA secondary structure G-quadruplexes are important for TRF2 binding to IL1R1 promoter. A. Schematic depicting G4 motifs (A and B) and their respective positions on the *IL1R1* promoter; sequence scheme depicting generic G4 motifs; B. Circular dichroism profile (220-310 nm wavelength) of oligonucleotides (5 µM) with sequence of the G4 motifs A or B (or respective mutants with base substitutions) in solution. C. ChIP-qPCR following BG4 ChIP at the IL1R1 promoter (fold enrichment over mock) overlaid with TRF2 occupancy (fold enrichment over IgG) in HT1080 cells. [N=3] Statistical significance was calculated using Šídák’s multiple comparisons test (p values: * ≤ 0.05, ** ≤ 0.01, *** ≤ 0.001, **** ≤ 0.0001). D. IL1R1 promoter activity in HT1080 cells from luciferase-reporter without or with G4-deforming substitutions against G4 motif A (IL1R1G4-MUT-A) or G4 motif B (IL1R1-G4 MUT-B) in presence or absence of TRF2 induction; G4 motifs A and B as shown in schematic; specific base substitutions have been indicated in blue font. [N=3] Statistical significance was calculated using Tukey’s multiple comparisons test (p values: * ≤ 0.05, ** ≤ 0.01, *** ≤ 0.001, **** ≤ 0.0001). E. *IL1R1* promoter activity, mRNA and protein levels in presence/absence of the G4 binding ligand 360A with or without TRF2 induction. For Left and centre panels-[N=3] Statistical significance was calculated using Tukey’s multiple comparisons test (p values: * ≤ 0.05, ** ≤ 0.01, *** ≤ 0.001, **** ≤ 0.0001). F. IL1R1 promoter-firefly luciferase reporter cassette, with or without substitutions deforming the G4 motif B, was artificially inserted at the CCR5 locus using CRISPR-cas9 gene editing in HEK293T cells (scheme in left panel). Promoter activity from (right panel). [N=3] Statistical significance was calculated using Tukey’s multiple comparisons test (p values: * ≤ 0.05, ** ≤ 0.01, *** ≤ 0.001, **** ≤ 0.0001). G. TRF2 occupancy by ChIP-qPCR (right panel) at the artificially inserted IL1R1 promoter without or with G4-deforming substitutions (IL1R1G4 MUT-B);ChIP-qPCR primers designed specific to the inserted promoter using the homology arms( see methods). [N=3] Statistical significance was calculated using unpaired T test with Welch’s correction (p values: * ≤ 0.05, ** ≤ 0.01, *** ≤ 0.001, **** ≤ 0.0001). Error bars correspond to standard error of mean from independent experiments.

### TRF2-dependent recruitment of histone acetyltransferase p300 induces H3K27ac modification at the *IL1R1* promoter

We had previously reported that TRF2 occupancy alters the local epigenetics at gene promoters(Mukherjee et al., 2018). Here we sought to understand if TRF2 occupancy brought about any epigenetic alteration at the *IL1R1* promoter. Screening of activation (H3K4me3 and H3K27ac) and repressive (H3K27me3, H3K9me3) histone marks at the *IL1R1* promoter following TRF2 induction, showed enhanced H3K27acetylation (**Supplementary Figure 4A**). In both HT1080 and MDAMB-231 cells, *IL1R1* promoter H3K27 acetylation increased/reduced on TRF2 induction/silencing respectively (**Figure 4A**). We sought to find a potential histone acetyltransferase (HAT) and noted canonical HATs p300/CBP(Roth et al., 2003) at the *IL1R1* promoter in multiple cell lines in ENCODE (**Supplementary Figure 4B**). ChIP-qPCR confirmed p300 occupancy within upstream 200-400 bp promoter of *IL1R1* in TRF2-up conditions, which reduced significantly on TRF2 downregulation, in HT1080 and MDAMB-231 cells (**Figure 4B**). Furthermore, occupancy of acetylated-p300/CBP (activated HAT) was enriched at the *IL1R1* promoter on TRF2-induction in HT1080 cells (**Figure 4C**). Together these showed TRF2-mediated recruitment and acetylation of p300/CBP, resulting in H3K27 acetylation at the *IL1R1* promoter. *IL1R1* promoter occupancy of p300, CBP, ac-p300/CBP and H3K27ac was relatively low in HT1080-LT compared to HT1080 cells with shorter telomeres (**Figure 4D**) consistent with lower TRF2 occupancy at the *IL1R1* promoter in LT cells (**Figure 1A**).

**Figure 4:**
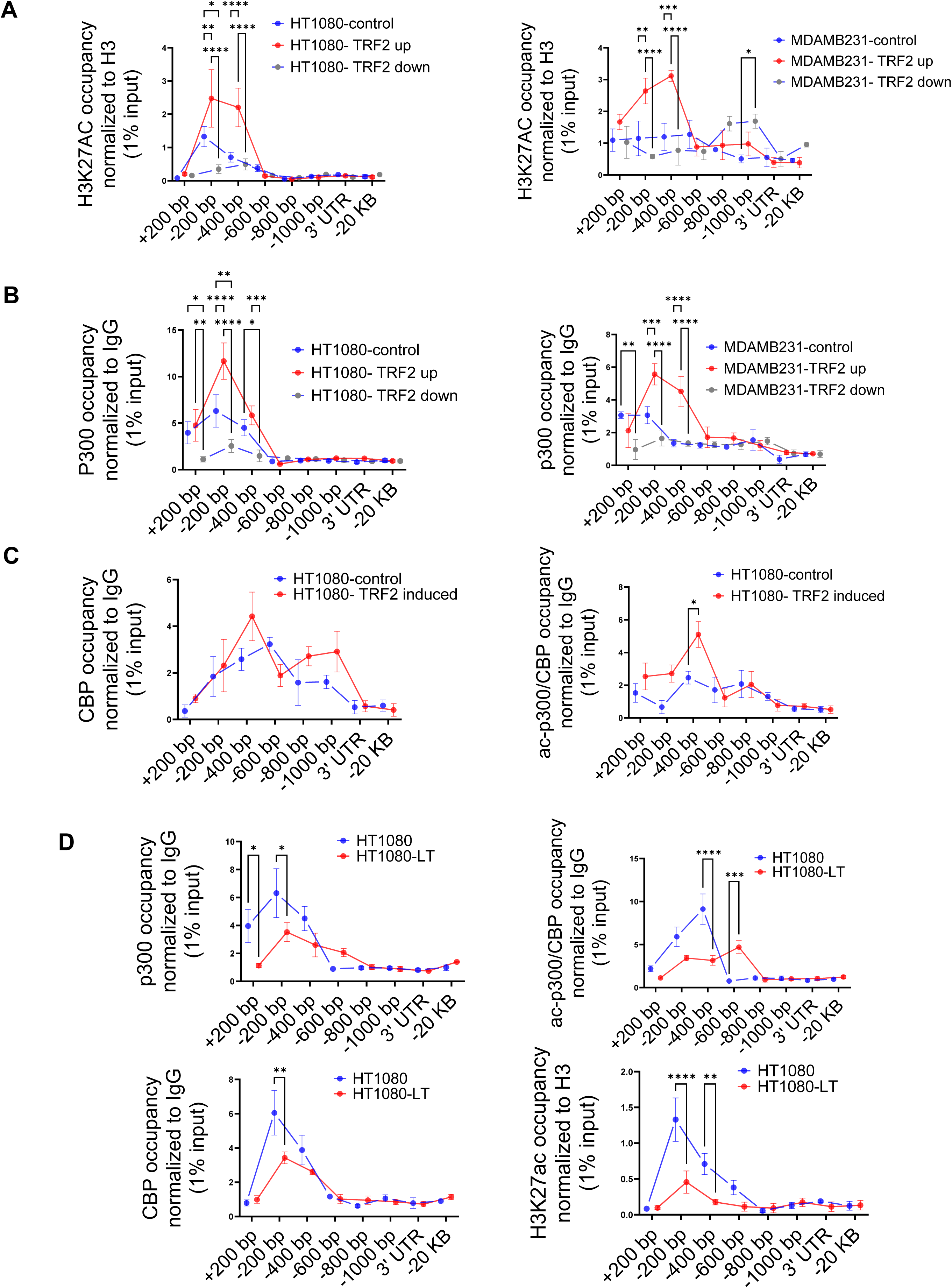
TRF2 recruits the histone acetyl transferase p300 to the *IL1R1* promoter. A.H3K27ac occupancy at the *IL1R1* promoter spanning +200 to -1000 bp of TSS by ChIP-qPCR in HT1080 (left panel) and MDAMB231 cells (right panel) in control (uninduced), TRF2-induced (up) or TRF2-down conditions; IL1R1-3’UTR or a region 20 kb upstream were used as negative controls. [N=3] Statistical significance was calculated using Tukey’s multiple comparisons test (p values: * ≤ 0.05, ** ≤ 0.01, *** ≤ 0.001, **** ≤ 0.0001). A. p300 occupancy on the *IL1R1* promoter in HT1080 (left panel) and MDAMB231 cells (right panel) in control (uninduced), TRF2-induced or TRF2-down conditions; IL1R1-3’UTR or a region 20 kb upstream were used as negative controls. [N=3] Statistical significance was calculated using Tukey’s multiple comparisons test (p values: * ≤ 0.05, ** ≤ 0.01, *** ≤ 0.001, **** ≤ 0.0001). B. CBP (left panel) and ac-p300/CBP (right panel) occupancy on the *IL1R1* promoter in HT1080 cells in control (uninduced) or TRF2-induced conditions; IL1R1-3’UTR or a region 20 kb upstream were used as negative controls. [N=3] Statistical significance was calculated using Šídák’s multiple comparisons test (p values: * ≤ 0.05, ** ≤ 0.01, *** ≤ 0.001, **** ≤ 0.0001). C. p300, CBP, acp300/CBP and H3K27Ac occupancy on the *IL1R1* promoter in HT1080 and HT1080-LT cells; IL1R1-3’UTR or a region 20 kb upstream were used as negative controls. [N=3] Statistical significance was calculated using Šídák’s multiple comparisons test (p values: * ≤ 0.05, ** ≤ 0.01, *** ≤ 0.001, **** ≤ 0.0001). All protein ChIP, other than histones, were normalized to 1% input and fold-change have been calculated over respective IgG. Histone ChIP were normalized to 1% Input and fold change over total H3 have been calculated for individual samples (Further details in the methods section) Error bars correspond to standard error of mean from independent experiments.

A recent work showed that TRF2 can physically interact with p300 and p300 acetylates TRF2 at specific lysine residues(YR and IK, 2013). We found that TRF2 directly interacted with p300, evident from co-immunoprecipitation using lysate from HT1080 cells (**Supplementary Figure 4C**). To ascertain direct TRF2-p300 interaction and consequent H3K27 acetylation, we did histone-acetyl-transfer assays using purified p300, histone H3, and acetyl-CoA (substrate for the acetyl group) with or without recombinant TRF2. Transfer of the acetyl group to histone H3 was significantly enhanced in the presence of both TRF2 and p300, relative to p300 only; and TRF2 co-incubated with p300 without H3 gave marginal HAT activity (**Supplementary Figure 4D**). These together demonstrate a direct function of TRF2 in recruitment and HAT activity of p300 in H3K27 acetylation at the *IL1R1* promoter.

### Acetylation of TRF2 at Lysine residue 293 is necessary for p300 recruitment on the *IL1R1* promoter

We sought to understand if the acetylation of TRF2 was important for p300-TRF2 complex recruitment to the *IL1R1* promoter. We found that several residues on TRF2 have been reported to be acetylated /deacetylated (Rizzo et al., 2017; YR and IK, 2013) (**Figure 5A**). To test the direct function of TRF2 acetylation in p300 recruitment, Flag-TRF2 mutants (K176R, K179R, K190R and K293R (Rizzo et al., 2017; YR and IK, 2013)) were screened (**Figure 5B**) for effect on *IL1R1* expression. TRF2-K293R gave notably compromised activation of *IL1R1* at mRNA and protein levels (**Figure 5B-D**). The binding of flag-TRF2-293R at the *IL1R1* promoter was similar to TRF2-WT (**Figure 5E**) suggesting that the loss in *IL1R1* activation was not due to lower promoter binding by TRF2-293R. However, TRF2-293R significantly reduced p300 and ac-p300/CBP recruitment on the *IL1R1* promoter relative to TRF2-wildtype (WT) induction (**Figure 5E**). TRF2 293R also showed lower physical interaction with p300 in an immunoprecipitation experiment in comparison to TRF2 WT (**Supplementary Figure 5A**). The TRF2 mutant K293R, devoid of H3K27 acetylation activity, gave a loss of function (**Figure 5B-D**). Therefore, TRF2-K293R was fused to dead-Cas9 (dCas9-TRF2-K293R), and dCas9-TRF2 was used as control. On using sgRNA against the *IL1R1* promoter, we found significant activation or downregulation of *IL1R1* in the case of dCas9-TRF2 or dCas9-TRF2-K293R, respectively, in HT1080 and MDAMB231 cells. In contrast, other TRF2-target genes (Mukherjee et al., 2018) remained unaffected indicating single-gene specificity (**Figure 5F**). We further checked the role of the TRF2 mutant K293R on other targets of TRF2 that we had previously validated(Mukherjee et al., 2019a) (**Supplementary Figure 5B**). While the TRF2 mutant K293R did not show much effect on genes wherein TRF2 acted as a transcriptional repressor, the effect on the candidate ‘activation’ targets of TRF2 was not uniform as well. The Lysine 293 in TRF2 seems to be important for activation but not repression. However, while the mutant K293R diminished TRF2-mediated activation of *IL1R1* and *PDGFR-B*, it did not affect *WRNIP1*. This suggests that the TRF2 PTMs might have context-dependence at specific loci in the genome.

**Figure 5:**
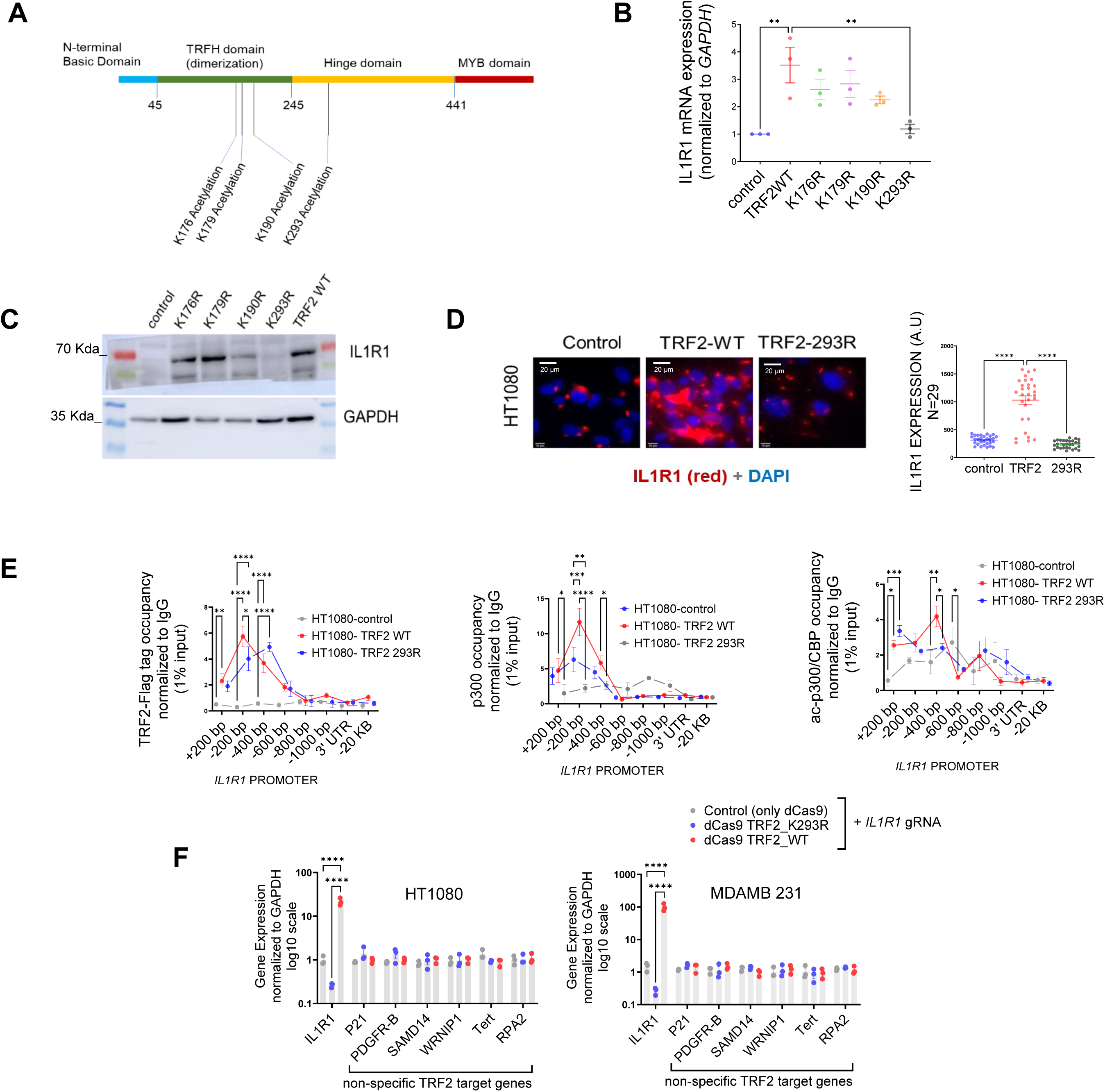
Acetylation of TRF2 at Lysine residue 293 is necessary for p300 recruitment on the *IL1R1* promoter. A. Schematic showing positions ( amino acid residues) where acetylation and deacetylation of TRF2 have been reported. B. IL1R1 mRNA expression post 48 hrs. of transient over-expression of various TRF2 acetylation mutants. *GAPDH* was used for normalization. [N=3] Statistical significance was calculated using Dunnett’s multiple comparisons test (p values: * ≤ 0.05, ** ≤ 0.01, *** ≤ 0.001, **** ≤ 0.0001). C. IL1R1 protein level following post 48 hrs. of transient over-expression of various TRF2 acetylation mutants; GAPDH used as loading control. D. Immunofluorescence for IL1R1 in HT1080 cells without (control) or with induction of flag-tag TRF2 WT or mutant TRF2-293R; quantification from 29 cells in each case shown in graph. [N=29] Statistical significance was calculated using Mann-Whitney’s non-parametric test (p values: * ≤ 0.05, ** ≤ 0.01, *** ≤ 0.001, **** ≤ 0.0001). E. Occupancy of flag-tag TRF2-WT or TRF2-293R on the IL1R1 promoter by ChIP-qPCR in HT1080 cells (left panel); IL1R1-3’UTR or a region 20 kb upstream were used as negative controls. Occupancy by ChIP-qPCR of p300 (middle panel) and acp300/CBP (right panel) without (control) or with induction of TRF2-WT and TRF2-293R. [N=3] Statistical significance was calculated using Tukey’s multiple comparisons test (p values: * ≤ 0.05, ** ≤ 0.01, *** ≤ 0.001, **** ≤ 0.0001). F. TRF2-wild type (WT) and TRF2-2293R mutant proteins fused with dCAS9 expressed and targeted to the IL1R1 promoter using *IL1R1*-specific gRNA in HT1080 and MDAMB231 cells. Following this, expression of *IL1R1* or other TRF2 target genes (non-specific with respect to the IL1R1-gRNA) in HT1080 (left) or MDMB231 cells (right). [N=3] Statistical significance was calculated using Tukey’s multiple comparisons test (p values: * ≤ 0.05, ** ≤ 0.01, *** ≤ 0.001, **** ≤ 0.0001). Error bars correspond to standard error of mean from independent experiments.

#### Cancer cells with higher TRF2 are sensitive to IL1-mediated NF-kappaB (p65) activation

Primarily ligands IL1A/B via receptors IL1R1/R2 activate IL1 signaling through NF-KappaB (p65/RELA)-Ser536 phosphorylation (A et al., 2010). This induces *IL1B* and other inflammatory genes including *IL6, IL8* and *TNF.* Here we checked NF-kappaB-Ser536 phosphorylation following stimulation with IL1B in the presence/absence of TRF2 in HT1080 and MDAMB-231 cells. TRF2-induction gave relatively enhanced NF-kappaB-Ser536 phosphorylation that sustained for a longer duration compared to uninduced controls in both cell types; total NF-kappaB levels remained unaltered (**Figure 6A-B**). Accordingly, on stimulation with either IL1A or IL1B in a TRF2-silenced condition, expression of the NF-kappaB targets *IL6, IL8* and *TNF* were reduced (**Figure 6C**). Further, the ligand TNFα was reported to induce NF-kappaB in an IL1-independent fashion (Liu et al., 2017) suggesting this pathway would be unaffected by the presence/absence of TRF2. Indeed, the expression of NF-kappaB targets (*IL6, IL8* and *TNF*) remained unaffected upon stimulation by TNFα in the absence of TRF2, supporting the role of TRF2 in IL1 signalling specifically (**Figure 6C**). Additionally, TRF2 induction enhanced the expression of *IL6*, *IL8* and *TNF*, which was rescued in the presence of the receptor antagonist IL1RA, further confirming the specific role of TRF2-IL1R1 in NF-kappaB activation (**Figure 6D**). When tested in NOD SCID mouse xenograft tumour samples, we observed that the ratio of phosphor-NF-kappaB to total NF-kappaB B was significantly higher in TRF2-induced tumours (**Figure 6E**).

**Figure 6:**
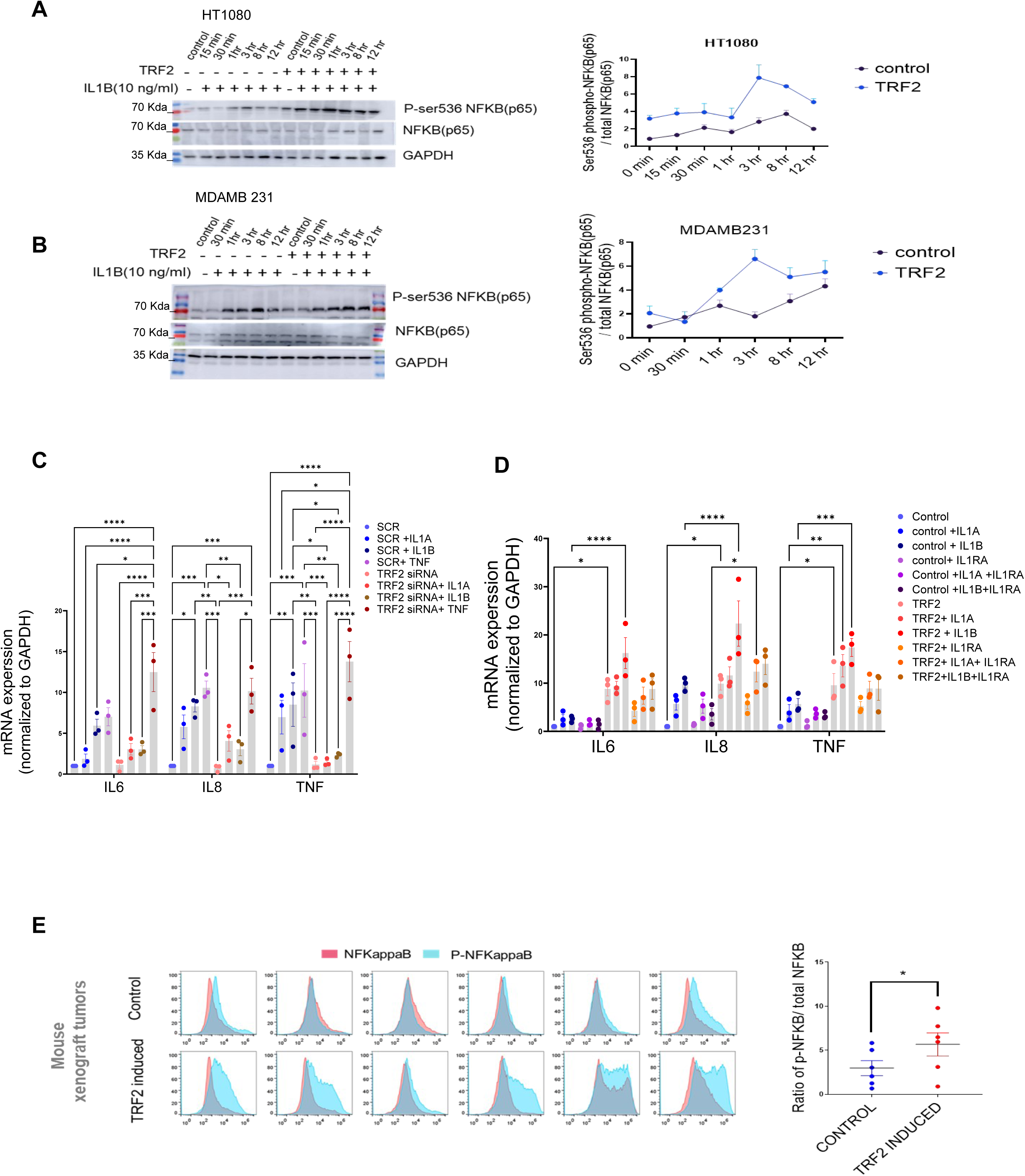
TRF2-dependent IL1 pathway activates NFkappa B (p65) in cancer cells. A-B. NFKappaB activation in presence of IL1B (10 ng/ml) in HT1080 (A) or MDAMB231 (B) cells with or without TRF2 induction; activation signaling was confirmed through NFKappaB-Ser536 phosphorylation (normalized to total NFKappaB); ratio of Ser566-p/total NFKappaB plotted for respective blots from three independent replicates (right panels). [N=2] C. Expression of NFKappaB targets *IL6, IL8* or *TNF* in presence/absence IL1A, IL1B or TNF-α (10 ng/ml) for 24 hr in control (scrambled siRNA) or TRF2-low (TRF2 siRNA) conditions in HT1080 cells. [N=3] Statistical significance was calculated using Tukey’s multiple comparisons test (p values: * ≤ 0.05, ** ≤ 0.01, *** ≤ 0.001, **** ≤ 0.0001). D. Expression of *IL6, IL8* or *TNF* in control or TRF2-induced conditions on treatment with either IL1A or IL1B (10 ng/ml) for 24 hr in absence (left panel) or presence (right panel) of the IL1-receptor-antagonist IL1RA (20 ng/ml) in HT1080 cells. [N=3] Statistical significance was calculated using Tukey’s multiple comparisons test (p values: * ≤ 0.05, ** ≤ 0.01, *** ≤ 0.001, **** ≤ 0.0001). E. NFkappaB/phosphor-Ser536-NFKappaB levels in xenograft tumors developed in NOD-SCID mice (control or doxycycline-induced TRF2 in HT1080 cells; N=6 mice in each group) by immuno-flow cytometry; mean fluorescence signal from individual tumors in control or TRF2-induced tumors plotted in adjacent graph; activation shown as ratio of pSer536-p-NFkappaB over total NFkappaB; significance was calculated using Wilcoxon’s non-parametric test. [N=6] Statistical significance was calculated using Mann-Whitney’s non-parametric test (p values: * ≤ 0.05, ** ≤ 0.01, *** ≤ 0.001, **** ≤ 0.0001). Error bars correspond to standard error of mean from independent experiments.

As shown in **Figure 5F** (mRNA expression of *IL1R1*), dCas9-TRF2-K293R diminished IL1R1 levels (**Supplementary Figure 6A**).This resulted in reduced NF-kappaB-Ser536 phosphorylation indicating attenuated NF-kappaB activation in HT1080 and MDAMB 231 cells; dCas9-TRF2 as expected showed activation of IL1R1 and NF-kappaB (**Supplementary Figure 6A**).

To directly test the function of IL1R1 in TRF2-mediated NF-kappaB activation, *IL1R1*-knockout (KO; CRISPR-mediated) was made in HT1080 cells having doxycline-inducible-TRF2: NF-kappaB activation (Ser536-phosphorylation), including upregulation of the NF-kappaB-target genes *IL2, IL6* and *TNF*, was significantly reduced following TRF2 induction in the KO cells compared to HT1080 cells (**Supplementary Figure 6B**). Together, these demonstrate the direct role of TRF2 in the induction of IL1 signalling through upregulation of *IL1R1*.

We reasoned that in cells with longer telomeres IL1 signaling would be affected due to reduced TRF2 binding at the *IL1R1* promoter resulting in attenuated activation of *IL1R1*. In the case of HT1080-LT cells, notable increase in NF-kappaB-Ser536 phosphorylation, following IL1B stimulation, was notably delayed (3 hr in HT10180-LT vs 30 min in HT1080 respectively; **Supplementary Figure 6C**).

### IL1R1 expression in Triple Negative Breast cancer (TNBC) is sensitive to variation in telomere length

To check if our findings were clinically relevant, we used TNBC tumour samples to characterize telomere length and *IL1R1* expression. The IL1( interleukin 1) pathway is relevant in TNBC prognosis as indicated by past studies(Lappano et al., 2020; M et al., 2016), and therefore it was relevant for us to use TNBC samples. We checked 94 TNBC samples and variation in telomeres across tumours was evident (**Figure 7A-C**).

**Figure 7:**
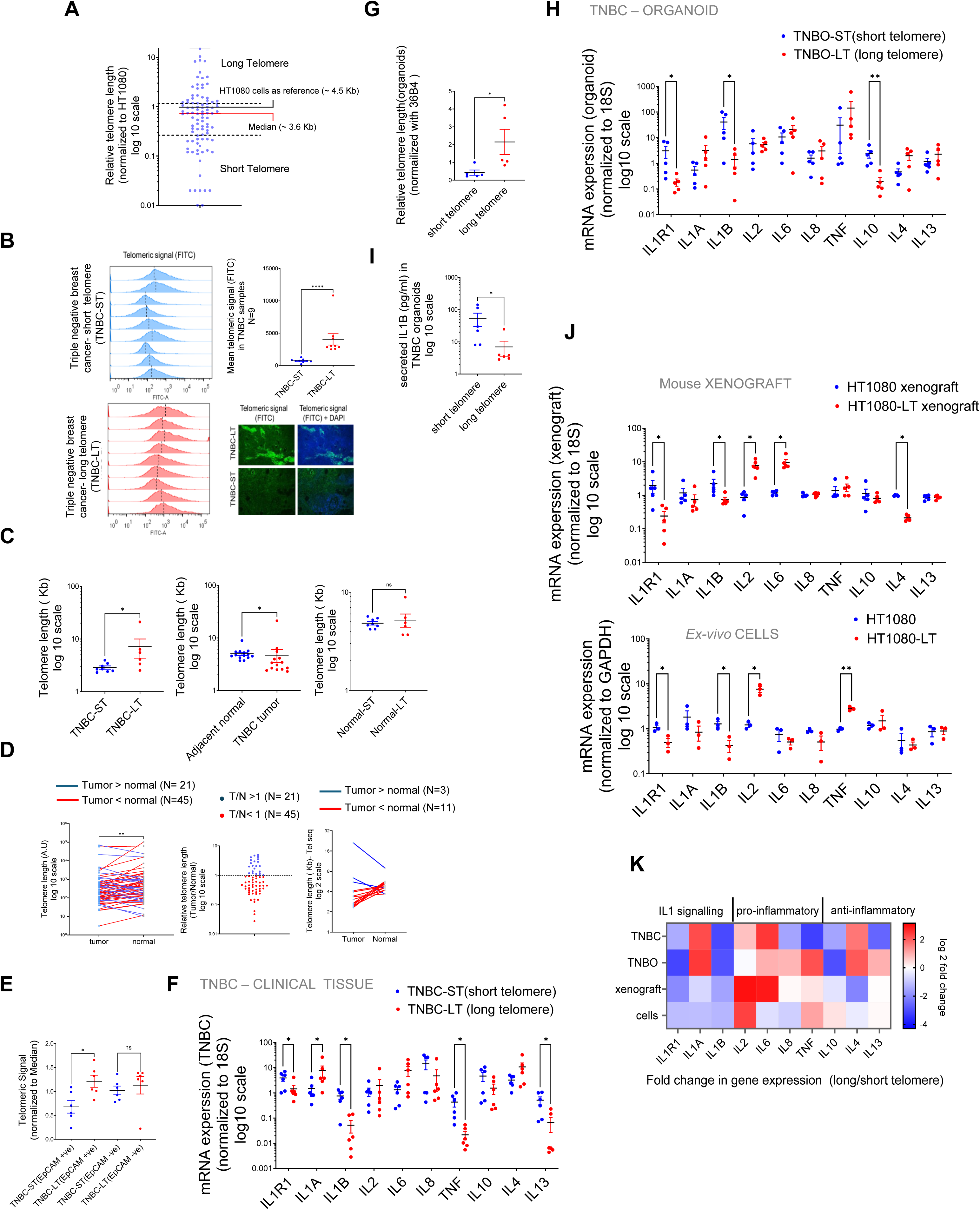
IL1R1 expression in Triple Negative Breast cancer (TNBC) samples is sensitive to inter-tumoral variation in telomere length. A. Relative telomere length of Triple Negative Breast Cancer (TNBC) samples (94 patients) by Telo-qPCR as reported earlier; signal from telomere-specific primers was normalized to single copy number gene 36B4 for individual samples. All samples were run with HT1080 DNA as control and telomeric signal from HT1080 cells (telomere length ∼4.5 Kb) was used as reference for relative measurement; median telomere length (3.6 Kb) shown by red bar; samples >50% or <50% of median (dotted lines) were designated as long or short telomere samples respectively. [ N=94] B. Flow cytometry analysis of telomere length of TNBC tissues from telomeric signal using telomere–specific FITC-labeled PNA probe; quantification of mean telomeric signal for nine TNBC-ST (short telomere, top left panel) and nine TNBC-LT (long telomere bottom left panel) shown in top right panel. Median telomere length has been indicated by dotted lines ( top right). TNBC tissue slides hybridized with telomere-specific PNA probes and counter stained with DAPI. Representative images for long/short telomere TNBC tissue shown ( bottom right) [N=9] Statistical significance was calculated using Mann-Whitney’s non-parametric test (p values: * ≤ 0.05, ** ≤ 0.01, *** ≤ 0.001, **** ≤ 0.0001). C. Telomere length was determined using the previously published algorithm Tel-Seq from sequenced genomes of TNBC samples (N=8 for TNBC-ST (short telomere) and N=6 for TNBC-LT (long telomere); and adjacent normal tissue from same patient). Samples identified as long-telomere (TNBC-LT) or short-telomere (TNBC-ST) using telo-qPCR were significantly different in TL, and consistent with telo-qPCR annotation (left); TL of tumor samples was lower that adjacent normal tissue [N=15] (center); and TL from adjacent normal tissues in LT or ST samples did not vary significantly [N= 8; N=6](right) Statistical significance was calculated using Mann-Whitney’s non-parametric test (p values: * ≤ 0.05, ** ≤ 0.01, *** ≤ 0.001, **** ≤ 0.0001). D. Paired telomere length assessment in tumor and adjacent normal tissue for 66 patients. Telomere signal was normalized to 36B4 (single copy gene ) by qRT PCR and compared between tumor and adjacent normal samples. The data was plotted pairwise (left) with lines connecting the individual tumor sample to the respective adjacent normal. All samples where tumor telomere length was lower than normal were connected by red lines and the samples where tumor telomeres where higher were connected by blue lines. Relative telomere length(Tumor/Normal) was plotted (middle) with samples with T/N<1 in red and T/N >1 in blue. Out of the 66 samples, 45 samples had T/N < 1, suggesting that about 2/3 rd of the samples tested had shorter telomeres in the tumors. Analysis of whole genome sequences of 14 pairs of samples using tel-seq pipeline revealed that 11 cases showed lower tumor telomere length compared to their adjacent normal counterparts (right). Left panel-Statistical significance was computed by Wilcoxon matched-pairs signed rank test (p values: * ≤ 0.05, ** ≤ 0.01, *** ≤ 0.001, **** ≤ 0.0001). E. TNBC tissue (N=6 each for long or short telomere samples) stained with EpCAM (far red) and hybridized with telomere-specific PNA probe (FITC). Mean telomeric signal was plotted in total EpCAM^+ve^ or EpCAM^−ve^ cells. [N=6] Statistical significance was calculated using Mann-Whitney’s non-parametric test (p values: * ≤ 0.05, ** ≤ 0.01, *** ≤ 0.001, **** ≤ 0.0001). F. mRNA expression of *IL1R1* and other key cytokines: *IL1A, IL1B, IL2, IL6, IL8*, *TNF, IL10, IL4* and *IL13* in long or short telomere TNBC tissue,; the *18S* gene was used for normalization. [N=6] Statistical significance was calculated using Mann-Whitney’s non-parametric test (p values: * ≤ 0.05, ** ≤ 0.01, *** ≤ 0.001, **** ≤ 0.0001). G. Relative telomere length of triple negative breast tumor organoids (TNBO) derived from TNBC samples: five TNBO-ST (short telomere) and five TNBO-LT (long telomere) was estimated by telo-qRTPCR. [N=5] Statistical significance was calculated using Mann-Whitney’s non-parametric test (p values: * ≤ 0.05, ** ≤ 0.01, *** ≤ 0.001, **** ≤ 0.0001). H. mRNA expression of *IL1R1* and other key cytokines in triple negative breast tumor organoids (TNBO) derived from TNBC samples-TNBO-ST and TNBO-LT; the *18S* gene was used for normalization. [N=5] Statistical significance was calculated using Mann-Whitney’s non-parametric test (p values: * ≤ 0.05, ** ≤ 0.01, *** ≤ 0.001, **** ≤ 0.0001). I. Secreted IL1B (pg/ml) from organoids TNBO-ST or TNBO-LT by ELISA using media supernatant form organoid culture (see methods).[N=6] Statistical significance was calculated using Mann-Whitney’s non-parametric test (p values: * ≤ 0.05, ** ≤ 0.01, *** ≤ 0.001, **** ≤ 0.0001). J. mRNA expression of *IL1R1* and other key cytokines using xenograft tumors (made from either short (HT1080) or long telomere (HT1080-LT) fibrosarcoma cells, and *ex-vivo* HT1080 or HT1080-LT cells. The *18S* gene was used for normalization for the tumors ;for ex-vivo cells *GAPDH* was used for normalization. Top panel-[N=5] Statistical significance was calculated using Mann-Whitney’s non-parametric test (p values: * ≤ 0.05, ** ≤ 0.01, *** ≤ 0.001, **** ≤ 0.0001). ; bottom panel-[N=3] Statistical significance was calculated using unpaired T test with Welch’s correction (p values: * ≤ 0.05, ** ≤ 0.01, *** ≤ 0.001, **** ≤ 0.0001). K. Heat map summarizing the mRNA expression of key cytokines across the models shown above: TNBC tissue, TNBC derived organoids (TNBO), xenograft tumors or ex-vivo cells in long or short telomere cases. Fold change of gene expression (long with respect to short telomeres) color coded as per the reference legend along y-axis. Error bars correspond to standard error of mean from independent experiments.

We measured TL (telomere length) in tissues from 94 breast cancer [triple-negative breast cancer (TNBC)] patients: Median TL was ∼3.6 kb (quantitative-telomeric-RTPCR (O’Callaghan et al.) (Telo-qPCR-see methods); **Figure 7A, Supplementary file 3**). Tumours with TL <50% of median were designated TNBC-short-telomeres (TNBC-ST) and >50% of median length TNBC-long-telomeres (TNBC-LT) (**Figure 1D**). TLs were confirmed by flow cytometry and in-situ hybridization (**Figure 7B**).

Further, reads from the whole genome sequence of the TNBC samples were analyzed for estimation of TL using the reported pipline-TelSeq (Ding et al., 2014): TL difference between TNBC-ST and LT was clear, and cancer tissue had lower TL than adjacent normal tissue consistent with earlier reports(Aviv et al., 2017). We found that, while there were inter-tumour telomere length variations, about ∼ 70% or more (45 out of 66 TNBC samples and 11 out of 14 in tel-seq analysis) samples for which data was available for adjacent normal tissue – showed lower telomere length in tumour samples (**Figure 7D**).

EpCAM^+ve^ (tumour-cell marker) (Went et al., 2004) cells within the TME had significant heterogeneity in TL (lower in TNBC-ST vis-a-vis LT), whereas the difference in TL of EpCAM^-ve^ cells was insignificant indicating TL variation in TME was primarily from cancer cells (**Figure 7E** and **Supplementary Figure 7A**). Details for the Tel-seq data have been tabulated with details of TL estimation for individual samples (**Supplementary Figure 7B**). We surprisingly found that telomerase activity negatively correlated with telomere length in TNBC (**Supplementary Figure 7C**), and while *TERT* was not found to be significantly higher, *TERC* was found to be significantly enhances in TNBC-LT samples (**Supplementary Figure 7D**). This indicates that *TERC* is possibly a more important determinant for telomere length in TNBC.

Next, along with *IL1R1*, a curated set of pro- and anti-inflammatory cytokines commonly studied concerning tumour immunity (B Homey, 2002; Briukhovetska et al., 2021) was screened in TNBC tissue (12 patients-6 long telomere, 6-short telomere). We found that *IL1R1, IL1B, TNF and IL3* were lower in long telomere samples (**Figure 7F**). Patient-derived organoids (TNBC-organoids from short (TNBO-ST) or long (TNBO-LT) telomeres) (**Supplementary Figure 7E**) were made and tested for telomere length. We found that the expected difference in telomere length was observable between the TNBO-ST and TNBO-LT samples (**Figure 7G**). When the same set of genes as in **Figure 7F** was tested in the TNBO samples, the same trend was observed for *IL1R1*, *IL1B* and *TNF*. In this case, *IL10* was also lower in TNBO-LT samples (**Figure 7H**). Secreted IL1B into the organoid media was also found to be lower in TNBO-LT samples (**Figure 7I**). For comparison, the same genes were looked at in HT1080 tumour xenograft and ex-vivo cells with long/short telomeres (**Figure 7J**).

*IL1B* and *IL1R1* expression was significantly low in the case of tumours/cells with relatively long telomeres and notably consistent across clinical tissue, organoids, xenografts and cells, in contrast to the other cytokines which showed model-specific variation (**Figure 7K**). Additionally, mRNA expression correlation from 34 TNBC samples further confirmed the inverse (negative) correlation of *IL1R1* with RTL (relative telomere length) and the consistent positive correlation between *IL1R1* and *IL1B* (**Supplementary Figure 7F**); showed key IL1 signalling genes largely upregulated in the short-telomere cells. Interestingly, *TRF2* showed a strong positive correlation with *IL1R1* and *IL8* but not with other pro-inflammatory cytokines.

We, next, used publicly available TCGA gene expression data of breast cancer samples (BRCA) (**Supplementary file 4**) to assess the effect of *IL1R1* expression on cancer prognosis. We categorized samples based on *IL1R1* expression: *IL1R1* high (N=254) and *IL1R1* low samples (N= 709). Notably, overall patient survival was significantly lower in *IL1R1* high samples (Log-rank p value: 0.0149) (**Supplementary Figure 7G**). We also checked the frequency of occurrence of various breast cancer sub-types in *IL1R1* high and low samples (**Supplementary Figure 7H**). While invasive mixed mucinous carcinoma (the most abundant sub-type) was predominantly seen in *IL1R1* low samples, metaplastic breast cancer was only found within the *IL1R1* high samples. Interestingly, metaplastic breast cancer has been frequently found to be ‘triple negative’, i.e., ER-, PR- and HER2-. (Reddy et al., 2020).

### Telomere length sensitive IL1R1 expression modulates Tumour-associated macrophage (TAM) infiltration in TNBC

Previous studies have implicated many telomere-sensitive genes to be related to immune regulation (Barthel et al., 2017; Hirashima et al., 2013). A recent study analyzed transcriptomic data across 31 cancer types to identify genes that are altered with telomere length(Barthel et al., 2017). The authors found that while sub-telomeric genes (up to 10 MB from the nearest telomeres) were altered with changes in telomere length as previously ascribed to the Telomere position effect (TPE) (Baur et al., 2001; Kim et al., 2016; Pedram et al., 2006; Robin et al., 2015), many ‘telomere-sensitive’ genes were distributed far from telomeres throughout the genome. This is consistent with a previous report that telomeres can affect transcription in regions far beyond sub-telomeres independent of TPE(Mukherjee et al., 2018). An intriguing observation made by this study was that when the top 500 genes to be altered with telomere elongation were functionally segregated, the highest number of genes fell under the category of ‘Immune response’. When a Gene Ontology analysis was done by us using these genes, ‘macrophage inhibitory factor signalling’ was the top biological process (**Supplementary Figure 8A**). Recent studies have implicated IL1R1-related signalling in M2 tumour-associated macrophage (TAM) infiltration in tumours(Chen et al., 2023; Zhang et al., 2022). In our data, we had observed consistent change in *IL1R1* and *IL1B* in long versus short telomere tumours and also noted that IL1 signalling has been widely implicated with TAM infiltration in multiple cancers (Carmi et al., 2009; Chittezhath et al., 2014; Lappano et al., 2020; Mantovani et al., 2019).

When breast cancer transcriptomic data from TCGA was segregated into high and low *IL1R1* groups (**Supplementary file 4**). Within the differentially expressed genes between high and low *IL1R1* groups, we looked at key immune markers such as reported TAM markers (*CD163, MRC1,D86,CD80*), TAM chemo-attractants (*CCL2, CCL3,CCL5, CCL8,CXCL12*) (Unver, 2019) as well as reported tumor infiltrating lymphocyte markers (*CD3G, CD4,CD8A, FOXP3, ENTPD1, PDCD1*) (**Figure 8A, Supplementary file 5**). We found that *MRC1* (gene coding for CD206-M2 macrophage marker) was the most significantly upregulated marker in *IL1R1* high samples. Interestingly, *TERT* (often correlated/associated with high telomere length) (Greider, 1998; McNally et al., 2019) was found to be significantly downregulated in *IL1R1* high samples. This observation aligns with our finding that *IL1R1* was higher in short telomere samples in TNBC and TNBO.

**Figure 8:**
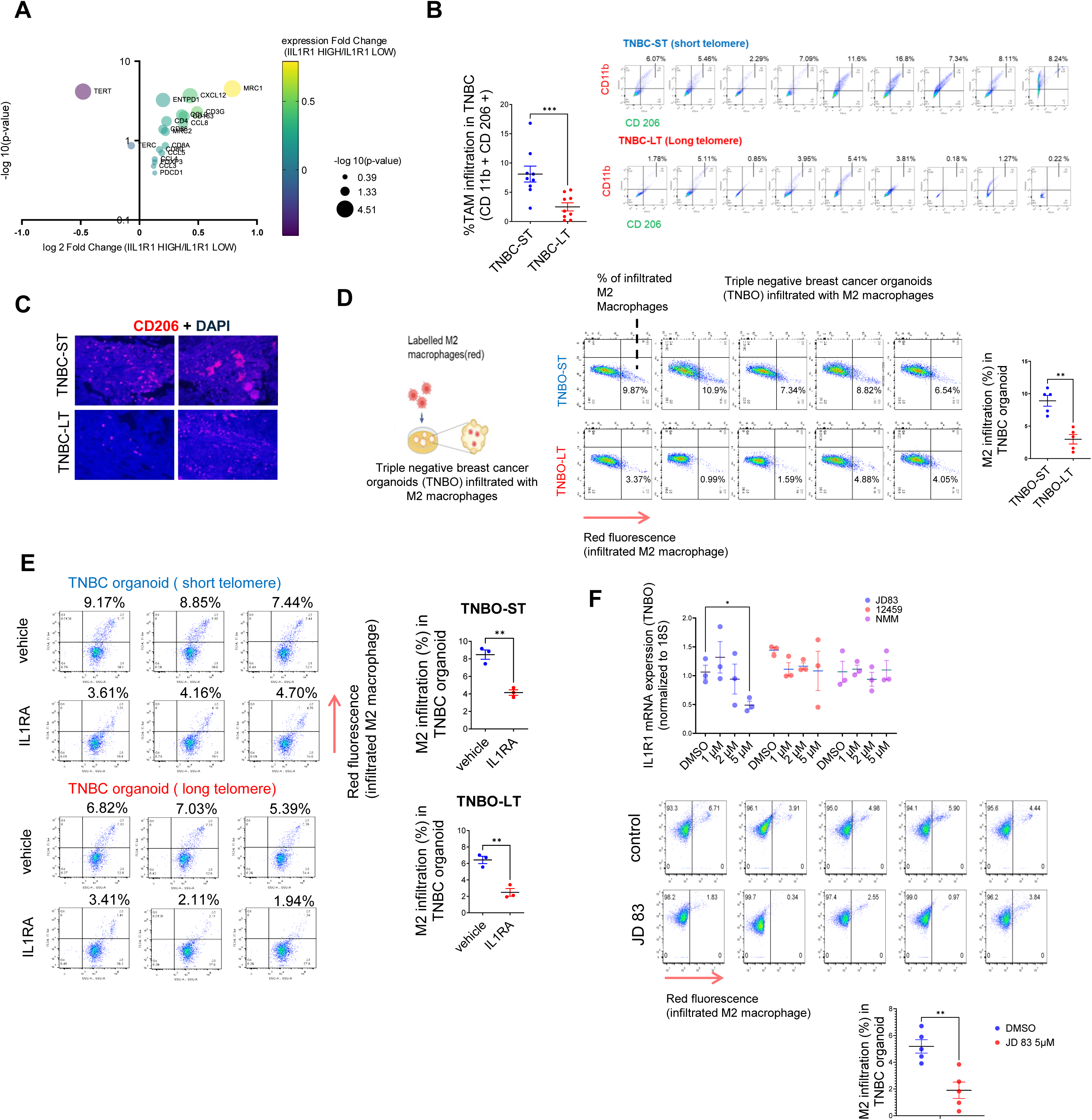
Telomere length sensitive IL1R1 expression modulates Tumor associated macrophage (TAM) infiltration in TNBC. A. Key Immune marker expression ( TAM markers, TAM related chemokines, T cell markers ), *TERT* and *TERC* were plotted for fold change of transcript level expression ( IL1R1 high / IL1R1 low) and adjusted p-values in breast cancer (BRCA) samples categorized as IL1R1 high or low from TCGA ( See supplementary file 4 ). B. Percentage tumor-associated-macrophage (TAM) infiltration in TNBC tissue with short (TNBC-ST) or long (TNBC-LT) telomeres using markers CD11b and CD206 (CD11b^+^CD206^+^ cells shown in top right quadrant); quantification of TAM infiltration from individual TNBC-ST or TNBC-LT samples have been plotted. [N=9] Statistical significance was calculated using Mann-Whitney’s non-parametric test (p values: * ≤ 0.05, ** ≤ 0.01, *** ≤ 0.001, **** ≤ 0.0001). C. TNBC tissue slides stained with TAM-specific marker CD206 for macrophage infiltration within tissue (counterstained with DAPI); representative images for two independent long or short telomere TNBC tissues shown. D. Labelled (red) M2 macrophages derived from THP1 cells incubated with triple negative breast tissue-derived organoids (TNBO) for 12 hours (scheme in left panel). Infiltration of M2 macrophages in organoids shown as percentage of labelled (red) M2 macrophages in individual flow-cytometry plots (florescence signal (FL3) in x-axis in log scale); percentage infiltration from five TNBO-ST or five TNBO-LT samples shown in right panel. [N=5] Statistical significance was calculated using Mann-Whitney’s non-parametric test (p values: * ≤ 0.05, ** ≤ 0.01, *** ≤ 0.001, **** ≤ 0.0001). E. Labelled (red) M2 macrophage infiltration in presence/absence of receptor antagonist IL1RA (20 ng/ml) for tumor organoids with short (TNBO-ST) or long telomeres (TNBO-LT) in three replicates; percentage values for M2 infiltration plotted in right panel. Red florescence (FL3; y-axis in log scale) and respective percentage M2 infiltration values marked on top of individual flow cytometry plots. [N=3] Statistical significance was calculated using unpaired T test with Welch’s correction (p values: * ≤ 0.05, ** ≤ 0.01, *** ≤ 0.001, **** ≤ 0.0001). F. mRNA expression of *IL1R1* was checked in presence of varying concentrations of G-quadruplex binding ligands ( 48 hr treatment) using TNBO. *18S* gene was used for normalization. Following this, the ligand JD83 was used to check M2 infiltration in TNBO as in (C-D). The M2 infiltration has been plotted for control and JD 83 treated samples. Top panel-[N=3] Statistical significance was calculated using unpaired T test with Welch’s correction (p values: * ≤ 0.05, ** ≤ 0.01, *** ≤ 0.001, **** ≤ 0.0001).; bottom panel-[N=5] Statistical significance was calculated using Mann-Whitney’s non-parametric test (p values: * ≤ 0.05, ** ≤ 0.01, *** ≤ 0.001, **** ≤ 0.0001). Error bars correspond to standard error of mean from independent experiments.

With these in mind, we checked TAM infiltration in TNBC-ST vis-à-vis TNBC-LT. Nine TNBC-ST and nine TNBC-LT tumours were analyzed for M2-type TAMs by double-positive CD11b (monocyte-derived cells) (Cassetta et al., 2016) and CD206 (M2-specific) (Jaynes et al., 2020) labelling: TNBC-LT had relatively low TAM infiltration (mean 1.78%, range: 0.18 – 5.41%) than TNBC-ST (mean 7.34%, range: 3.76-16.80%; **Figure 8B,Supplementary Figure 8B**). Accordingly, CD206 expression was found to be lower in tissue sections from TNBC-LT relative to TNBC-ST tumours (**Figure 8C**). We further found that the total proportion of immune cells (% of CD45 +ve) did not vary significantly between short and long telomere TNBC samples (**Supplementary Figure 8C**). However, TNBC-ST samples had a higher percentage of myeloid cells (CD11B +ve) within the CD 45 +ve immune cell population. We checked in three TNBC-ST and TNBC-LT samples and found that the percentage of M1 macrophages (CD86 ^high^ CD 206 ^low^) in the myeloid population was lower than that of the M2 macrophages (CD 206 ^high^ CD 86 ^low^) and unlike the latter, did not vary significantly between TNBC-ST and TNBC-LT samples (**Supplementary Figure 8C**).

We used the available data for *TERT/TERC* expression for a subset of the samples and plotted the telomerase activity for these TNBC samples (**Supplementary Figure 8D**). Upon checking we found that while *TERT* did not significantly correlate with either telomere length or telomerase activity, the correlation with *TERC* was significant with telomere length. *TERC* was earlier shown to vary significantly between short and long-telomere TNBC samples (**Supplementary Figure 7D**). Therefore, in TNBC, *TERC* seems to be critical for telomere length. Individually, neither *TERT* nor *TERC* correlated significantly with % TAM in TNBC tissue. As expected from our other results, % TAM correlated significantly with RTL. This indicates that in TNBC tissue, telomere length is possibly a more important determinant for TAM infiltration than either *TERT*, *TERC* or telomerase activity.

TAM infiltration was further tested using patient tumour-derived organoids from five long and five short telomere cases. M2-type macrophage derived from THP1 cells (**Supplementary Figure 8E**) were labelled (red-tracker dye) before co-incubating with each of the five TNBO-ST or five TNBO-LT: Following 12 hr incubation TNBO-ST had on average 8.56% infiltration of M2-type macrophages (range: 6.54-10.9%) compared to 3.14% (range: 0.99-4.88%) for TNBO-LT (**Figure 8D**).

Further, we used the IL1R1-receptor antagonist IL1RA to directly test IL1R1 dependence. The minimum dosage of IL1RA where expression of inflammatory markers was found to be altered in TNBO was used in the assays (**Supplementary Figure 8F**). Organoids treated with IL1RA before co-incubating with dye-labelled M2-type macrophages showed >50% reduction in macrophage infiltration (**Figure 8E**).

Finally, we screened several G4 binding ligands in MDAMB231 cells and found that a few ligands (JD83, 12459 and NMM) could reduce *IL1R1* expression (**Supplementary Figure 8G**). These ligands were tested in TNBO and we found JD83 to be effective in reducing *IL1R1* expression at 5 µM concentration (**Figure 8F**). We found that JD83 is also effective in reducing M2 migration in TNBO, showing promise for future therapeutic interventions.

## Discussion

Recently we described the TL-sensitive expression of genes distant from telomeres (Mukherjee et al., 2018). Cells with relatively long telomeres sequestered more TRF2 at the telomeres affecting non-telomeric binding, and vice-versa. Proposed as the telomere-sequestration-partition (TSP) model (Vinayagamurthy et al., 2020), this is independent of telomere-positioning-effect (TPE) on expression of sub-telomeric genes (up to 10 Mb from telomeres) (Robin et al., 2014). TL-dependent *IL1R1* activation by non-telomeric TRF2 is in tune with the TSP model. It needs to be noted that the *IL1R1* gene is located above 100 MB away from the nearest telomere in humans, and hence far out of the proposed 10MB range of the TPE model. Altered transcription when telomere maintenance is affected in varied contexts (Hirashima et al., 2013; Mukherjee et al., 2018), particularly across 31 cancer types(Barthel et al., 2017), appears consistent with TSP and support our observations related to TL-dependent IL1 signalling. A model of telomere length and TRF2-dependent *IL1R1* expression and the potential modulation of TAM infiltration is presented as a graphical summary (**Figure 9**).

**Figure 9.**
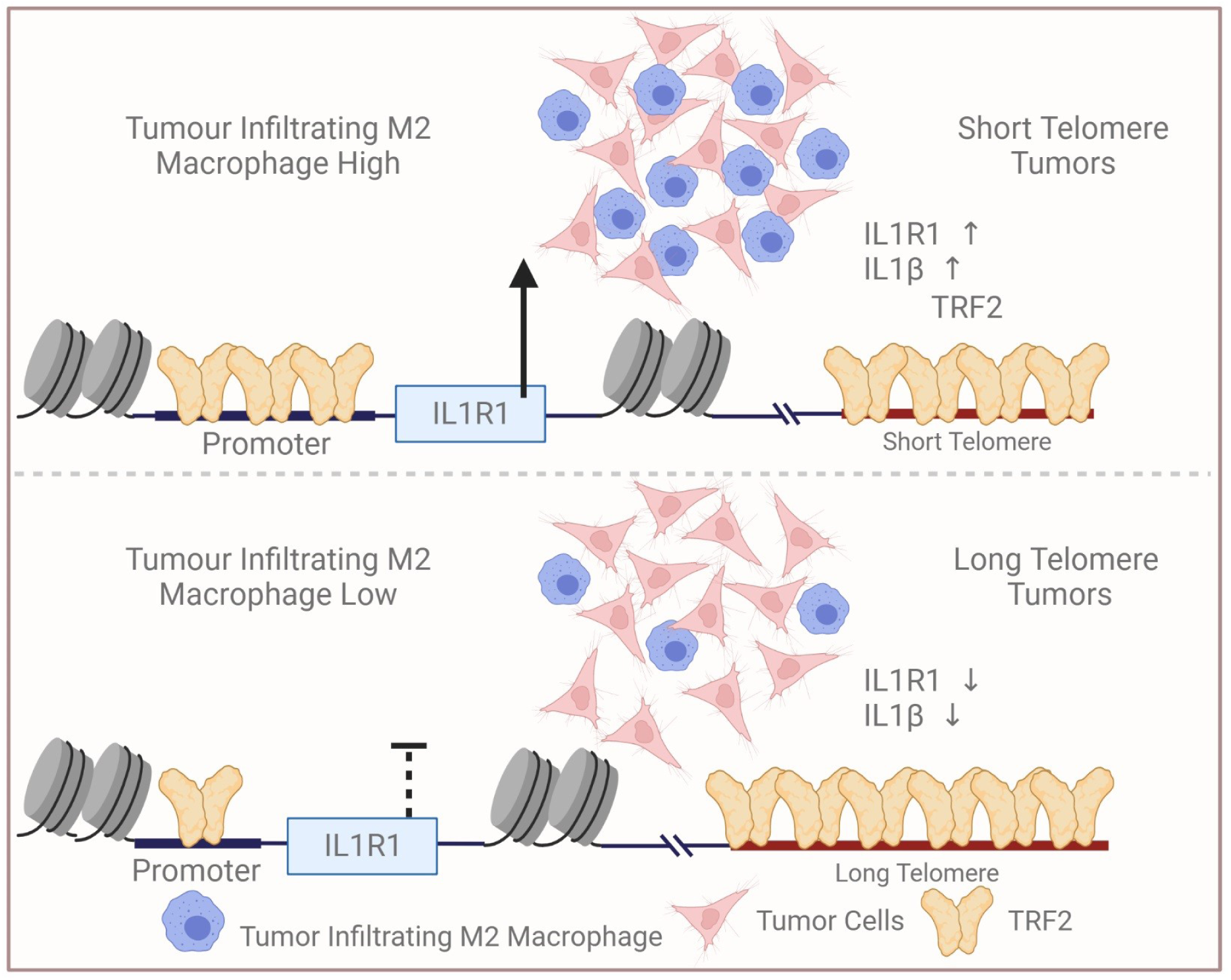
Scheme depicting relatively low infiltration of TAM in tumors with relatively long telomeres vis-à-vis tumors with short telomeres. Reduced non-telomeric TRF2 binding at the IL1R1 promoter in tumors with long telomeres, and consequent low IL1R1 activation, attenuated p65-mediated IL1-beta and macrophage infiltration. Figure created with BioRender.com.

Besides TRF2, the non-telomeric function of the shelterin protein RAP1 has been reported as part of complex with NF-kappaB (Teo et al., 2010). Although TRF2 has been implicated in influencing Natural Killer (NK) cells and infiltration of Myeloid Derived immuno-Suppressive Cells (MDSC) (Biroccio et al., 2013; Cherfils-Vicini et al., 2019) underlying mechanisms were unclear. However, the function of non-telomeric TRF2 in either IL1 signaling or in a telomere-sensitive role in immune signalling, has not been previously studied.

Mechanistically, we found that DNA secondary structure G4s present on the *IL1R1* promoter are necessary for TRF2 binding. Perturbation of the G4s genetically by destabilizing base substitutions, or upon using small molecules that bind to G4 inside cells resulted in reduced promoter TRF2 binding and *IL1R1* expression. This is in line with our earlier results showing G4-dependent non-telomeric TRF2 binding throughout the genome, and how TRF2 recruitment to specific promoters leads to epigenetic histone modifications (Mukherjee et al., 2019a, 2019b, 2018; Shalu Sharma et al., 2021). We recently noted the transcription repression of telomerase (*hTERT*) to be TRF2-mediated in a PRC2-repressor-dependent fashion(Shalu Sharma et al., 2021).This, interestingly, suggests the emerging role of TRF2 as a transcriptional modulator with repressor/activator (as shown here) functions depends on cofactor engagement that is possibly contextual.

The histone-modifying enzyme complex-p300/CBP have been keenly studied in cancer epigenetics specifically breast cancers (Iyer et al., 2004; Ramadan et al., 2021; Ring et al., 2020). Recent findings also show acetylated H3K27 enrichment at the telomeres through p300-dependent HAT activity (Cubiles et al., 2018); however role of TRF2 in p300 engagement or H3K27 acetylation at the telomeres was not investigated. Here, notably, we found TRF2-dependent p300 recruitment, and HAT activity resulting in H3K27 acetylation at the *IL1R1* promoter. Together, these support the role of non-telomeric TRF2 as an activating transcription factor through p300 implicating a broader role of TRF2-p300-dependent histone modifications in cancer. Further, these suggest the possibility of similar functions of telomeric TRF2 in H3K27 acetylation at the telomeres (Cubiles et al., 2018). The lysine residue K293 acetylation on TRF2 seems to be crucial for p300-TRF2 interaction.

It has been noted previously that TERT can have effects on NF-kappa B signalling independent of telomere length alterations(Ghosh et al., 2012). We observed that in long telomere HT1080-LT cells, while NF-kappa B signalling was retarded-a late surge of NFKappa-B activation occurred. We are inclined to think this could be a potential effect of TERT over-expression. Understanding the extent to which TERT influences the inflammatory signalling in TNBC independent of telomere length would require further work.

IL1 signalling in relation to inflammation in cancer has been extensively studied and consequently, IL1B blockers and the receptor antagonist IL1RA are of clinical interest (B Homey, 2002; Chittezhath et al., 2014; F Balkwill et al., 2001; Kaplanov et al., 2019; Lappano et al., 2020). Further, the effect of altered IL1 signalling on M2-type TAM infiltration was reported in multiple cancer types (Chanmee et al., 2014; Hu et al., 2015). TAM infiltration has been consistently linked to tumour aggressiveness (Oner et al., 2020) and recent work on targeting (Lee et al., 2019; TL and I, 2011) and modifying TAMs from M2-like states to a more anti-tumourigenic M1-like states (Jaynes et al., 2020) have shown promise as strategies for intervention in cancer. The clinical significance of blocking IL1 signalling in cancers has been widely reported (Briukhovetska et al., 2021; Kaplanov et al., 2019; Mantovani et al., 2018). Based on our findings, we attempted TRF2-mediated single-gene-targeted epigenetic silencing of *IL1R1* in cell lines and ligand-based approaches in TNBC organoids. A combination of single gene editing and small molecule/ ligand-based intervention might be useful in therapy.

For the functional significance of TL in TME, we focused on IL1-signalling and TAM. Results from TNBC organoids here show that blocking IL1R1 using IL1RA was sufficient for restricting M2 infiltration (**Figure 8E**) suggesting that the IL1 signaling within the tumour cell is important. However, particularly because macrophages are known to be a rich source of IL1B (Carmi et al., 2009; Duque and Descoteaux, 2014; Mantovani et al., 2018), and given the model-specific changes in inflammatory cytokines noted by us, further experiments will be required to understand how telomeres influence paracrine/ juxtracrine IL1 signalling between cancer cells and TAM.

It is quite possible that TL affects tumour-immune signalling through other immune cell types as well (implied by altered immune-response genes in TL-dependent gene-expression analysis (Barthel et al., 2017; Hirashima et al., 2013)) and/or influences the nature of TAM (Hung et al., 2016), which is known to vary contextually (Mantovani and Locati, 2013). These possibilities, together with the well-established effect of IL1-targeting in improving cancer prognosis (Briukhovetska et al., 2021; Cristofari and Lingner, 2006; Mantovani et al., 2019, 2018) underline the need for further work to attain a deeper understanding of telomere-sensitive IL1 signalling in the TME.

Although we observed a range of telomere lengths across cancers in TNBC patients (**Figure 7A-C**), as also noted by other groups (Londoño-Vallejo, 2004; Pellatt et al., 2013; Shen et al., 2007; Wentzensen et al., 2011; Willeit et al., 2010), tumours largely have been reported to maintain shorter telomeres than normal tissue (Aviv et al., 2017). Here we found that tumour cells with shorter TL-activated IL1 signalling increase M2-type TAM infiltration, which is known to aid immunosuppression (Allavena et al., 2008). Based on our results, and other reports suggesting immunomodulation in cancers with relatively short telomeres (Barthel et al., 2017; Hirashima et al., 2013), it is therefore tempting to speculate that short-telomere cancer cells might be ‘selectively’ better equipped to evade host immune cells. To sum up-our work reveals new molecular connections between telomeres and tumour immunosuppression, presenting a conceptual framework for a better understanding of patient-specific responses to cancer immunotherapy.

## Materials and Methods

### Cell lines, media and culture conditions

HT1080 fibrosarcoma cell line was purchased from ATCC. HT1080-LT cells(Cristofari and Lingner, 2006) were a kind gift from Dr. J. Ligner. HT1080 and MRC5 (purchased from ATCC) cells and corresponding telomere elongated cells were maintained in Modified Eagle’s medium (MEM) (Sigma-Aldrich) supplemented with 10% Fetal Bovine Serum (FBS) (Gibco). MDAMB231 (gift from Mayo Clinic, Minnesota, USA) and HEK293T (purchased from NCCS, Pune) cells were maintained in DMEM-HG (Dulbecco’s Modified Eagle’s medium - High Glucose) (Sigma-Aldrich) with a supplement of 10% FBS. All THP1 cell lines (gift from Dr Vivek Rao, IGIB, Delhi and subsequently purchased from NCCS, Pune) and derivative cell types were maintained in RPMI with 10% FBS and 1X Anti-Anti (anti-biotic and anti-mycotic from Thermo Fisher Scientific). All cultures were grown in incubators maintained at 37°C with 5% CO_2_. All stable lines generated from these cell types were maintained in the same media with 1X Anti-Anti.

### Processing of tumour/xenograft tissue

Tissues collected for this study were stored in RNA later (Sigma-Aldrich) at -80 °C. Approximately 10 milligrams of tumour tissue were weighed and washed thrice with ice-cold 1X PBS-1 ml (filtered). About 300 microlitres of PBS was left to submerge the tissue after the final wash. Using a sanitized razor blade, the tumour was cut into small pieces (the efficiency of the digestion is dependent on how well the tumour is minced; large pieces will not digest properly and may need extra digestion time). The tumour soup was transferred into a 15 ml tube using a P1000 pipette (cut the top of the tip to be able to take all the tissue) and the volume was adjusted to 4 ml of 1X pre-chilled PBS. 80 µl of Liberase TL stock solution (Roche) was mixed using a vortex and incubated for 45 min at 37 °C under continuous rotation. The sample was checked visually; the cell suspension should look smooth (if the cell suspension still contains tissue fragments digest for an additional 15 min). To end the enzymatic digestion,10 ml PBS with 1 % w/v BSA (Sigma-Aldrich) was added. The cells were filtered using a 100 µm cell strainer in a 50 ml tube and the volume adjusted to 20 ml with 1X PBS (pre-chilled). Cells were centrifuged for 7 min at 500 g and 4 °C. The supernatant was discarded and the cells were re-suspended in 1 ml Serum-free media. Subsequently, cells were counted and volume was made up to get a cell density of 1 million cells/ ml. Following this, the cell suspension was used for ChIP, DNA/RNA isolation and flow cytometry.

### Organoid generation and culture

TNBC tissue was used to make tumour organoids using the following protocol-

1. The fresh tissue was stored in 5 ml of Media 1 (Advanced DMEM F12 + 1X Glutamax + 10mM HEPES + 1X AA).
2. 1-3 mm^3^ tissue piece was cut, washed and minced in 10ml (Media 1).
3. The minced tissue was digested overnight on a shaker in Media 1 with 1-2mg/ml of collagenase.
4. After overnight incubation the digested tissue was sequentially sheared using a 1 ml pipette.
5. After each shearing step, the suspension was strained using a 100µm filter to retain tissue pieces. Thus entering in subsequent shearing step with approximately 10 ml Advanced DMEM F12.
6. To this, 2% FCS was added and centrifuged at 400 g.
7. The pellet was resuspended in 10mg/ml cold cultures growth factor reduced BME Type 2 and 40µl drops of BME suspension was allowed to solidify on pre-warmed 24 well culture plates at 37°C 5% CO_2_ for 30 min.
8. Upon complete gelation 400 µl of BC organoid medium was added to each well and plates transferred to 37°C 5% CO_2._

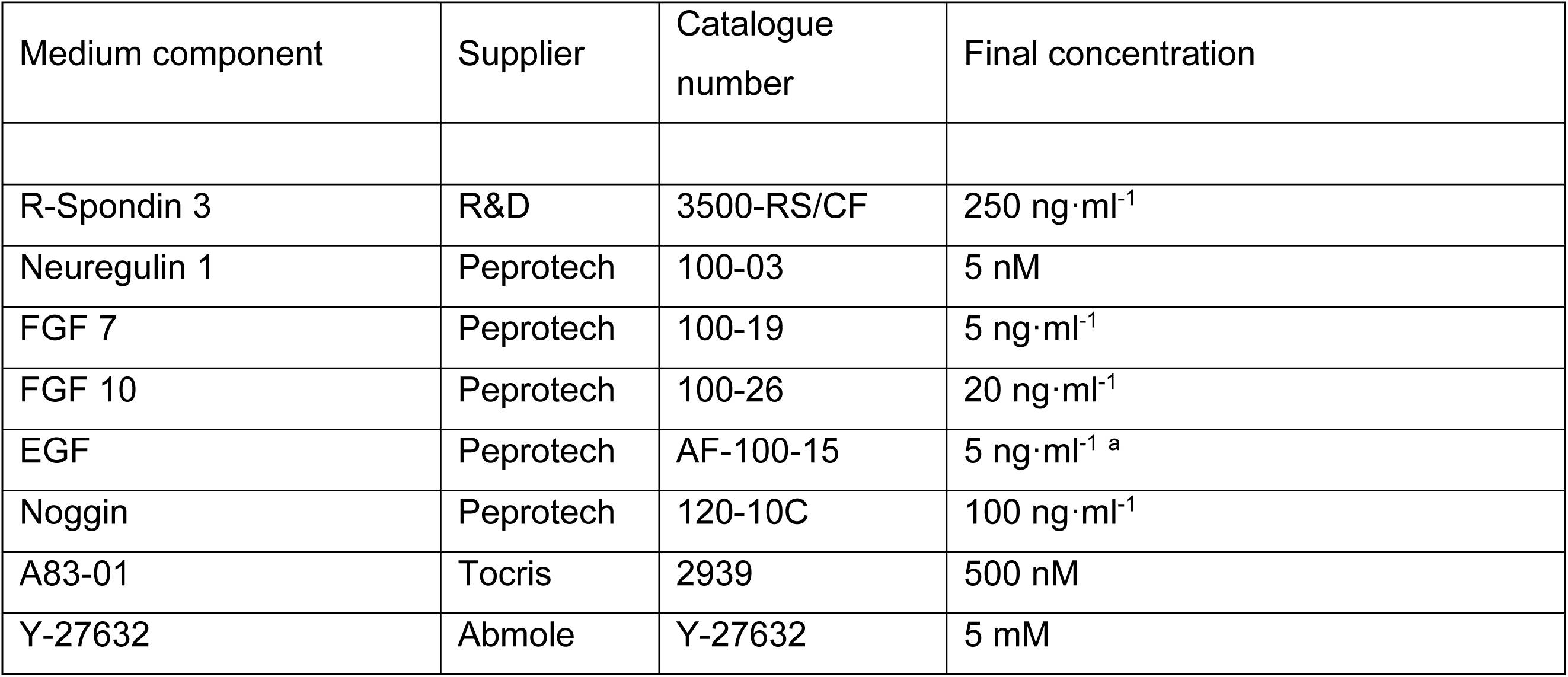
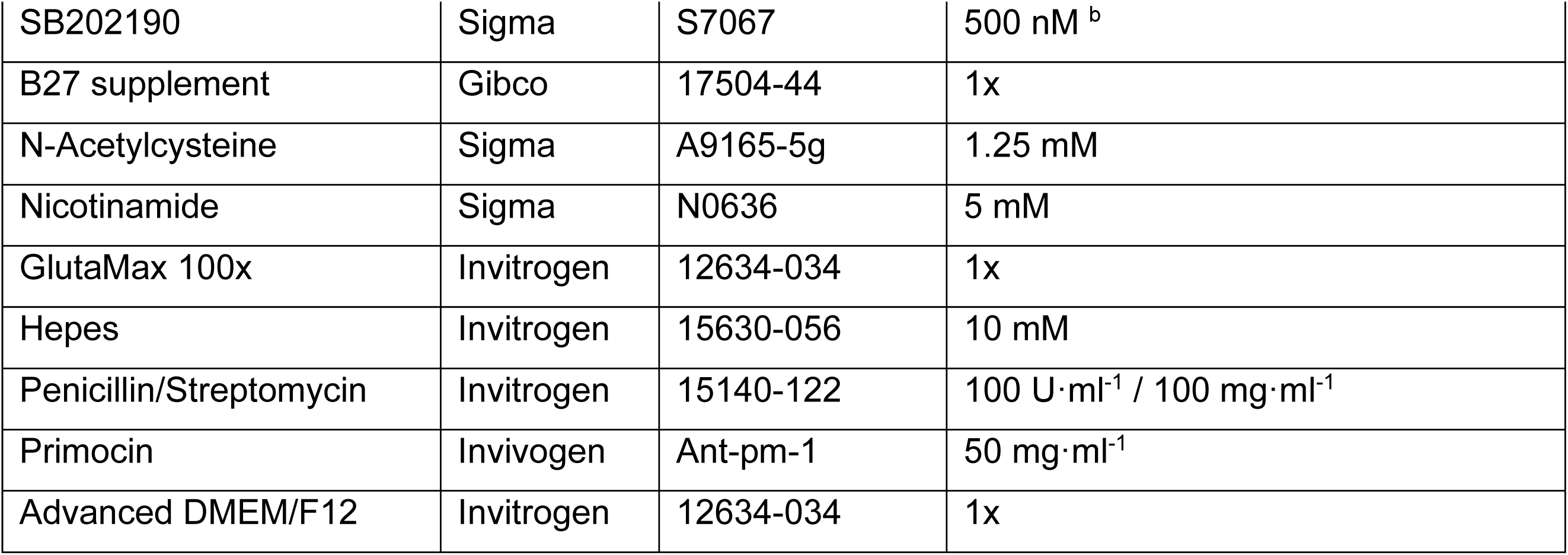
9. The media was changed every fourth day.
10. The culture was grown for 10 days before passaging.

II. Passaging Organoid Cultures

1. Cultrex growth factor reduced BME Type 2 geltrex was thawed overnight on ice at 4 °C.
2. Media was removed from the organoid wells and washed twice with PBS-EDTA.
3. For each well of 24 well plate 300 µl of TrypLE was added and incubated at 37°C 5% CO_2_ for 5 min.
4. 600 µl of washing media (DMEM F12) was added and organoids were dissociated vigorously by pipetting.
5. . Organoid solution was transferred to 15 ml falcon with 5 ml washing media and centrifuged at 1000rpm for 5 min to pellet organoids.
6. Supernatant was carefully aspirated leaving behind 200ul media and 500ul of fresh media, mixed well gently and transferred to 1.5ml tube for centrifugation at 650*g for 5min at 4 °C.
7. Supernatant was gently aspirated and resuspended in cold cultrex and 40ul drop/well was seed in a pre-warmed 24 well plate.
8. Incubated for 30 min and add 400 µl of BC organoid medium.

### M2 macrophage generation from THP1 cells

THP1 cells (0.5 million cells in 1ml) were seeded in a 35 mm dish and treated with PMA (10 ng/ml) (Sigma-Aldrich) for 24 hrs in RPMI complete media (+ 10% FBS) (Gibco). PMA was removed and cells were cultured for 48 hrs in RPMI to get semi-adherent M0 cells. Following this, cells were treated with IL4 (10 ng/ml) (Gibco) and IL13 (20 ng/ml) (Gibco) for 24 hrs in RPMI complete media. Cells were checked for expected morphological changes and trypsinized for use as M2 macrophages.

### M2 macrophage infiltration assay in TNBC tumour organoids

M2 macrophages were labelled with Cell Tracker red dye (Thermo Fisher Scientific) for 30 mins and the excess dye was washed off. 10000 labelled M2 macrophages in 20 ul 1X PBS were introduced in a 96 well with TNBC organoids in a dome with supplement enriched media (200 µl). 12 hrs. later, the media was washed off with 3 1X PBS washes and the organoid dome was extracted with 100 µl TrypLE (Gibco) and resuspended with 100 ul chilled 1X PBS. This cocktail was kept on ice for 5 mins and centrifuged at 1000g for 10 mins. The supernatant was removed and the process was repeated.

The cells were resuspended in 500 ul chilled 1X PBS and subjected to Flow-cytometric analysis for scoring percentage of labelled (red) M2 macrophages infiltrating the organoids.

### Xenograft tumour generation in NOD-SCID mice

The xenograft tumour generation in NOD SCID tumours was outsourced to a service provider ( Vivo Bio TechLtd – Telangana, India). 2.5 million cells (HT1080, HT1080-LT or HT1080-TRF2 inducible) were subcutaneously injected in healthy male mice (4-6 weeks old) and tumour growth was allowed. In case of TRF2 inducible HT1080 cells, tumours were allowed to grow to 100 mm^3^ before oral dosage of doxycycline (or placebo control) was started at 2 mg/kg of body weight of mice. Mice were sacrificed to obtain tumours when tumours reached an average volume of 600 mm^3^(± 100).

### Telomeric Fluorescent *in-situ* hybridization (FISH)

Telomere-specific PNA probe [TelC-FITC-F1009-(CCCTAA)n from PNABio] was used following the manufacturer protocol with minor modifications. Briefly, the cells grown in chamber slides up to ∼70% confluency and were treated with RNase solution for 5-10 mins in 1X PBS at room temperature. For tissue section, rehydrated tissue sections on slides were used prior to RNase treatment. Slides were washed in chilled PBS twice followed by dehydration with an increasing concentration of ethanol gradient. Slides were briefly warmed at 80°C and then 250 nM PNA was added with hybridization buffer (20 mM Tris, pH 7.4, 60% formamide (Sigma-Aldich), 0.5% of 10% Goat Serum). Slides were heated and covered with the buffer at 85°C for 10 mins, kept at room temperature in a dark humid chamber for 4 hrs to overnight to facilitate slow cooling and hybridization. The unbound PNA was washed with wash buffer (2X SSC, 0.1% Tween-20(Thermo Fisher Scientific)) followed by PBS wash twice. DAPI solution (Sigma-Aldrich) was added for 5 mins and then the slides were washed in PBS. ProLong Gold Antifade Mountant (Thermo Fisher Scientific) was added and the section was covered with a cover glass. Imaging was done with a Leica SP8 confocal microscopy system. Images were analyzed using open source image analysis software Fiji (Image J, imagej.net/software/fiji/).

### DNA and RNA extraction from TNBC/xenograft tissue

Allprep DNA/RNA Universal kit (Qiagen) was used as per manufacturer’s suggested protocol for extraction of DNA and RNA from single cell suspension prepared from tissues as described before.

### Telo-qPCR

For telomere length assessment by qRT PCR, telomere specific primers and single copy number gene 36B4 for normalization was used in PCR reactions with sample DNA using reported protocol(O’Callaghan et al.).

The primers used are as follows:

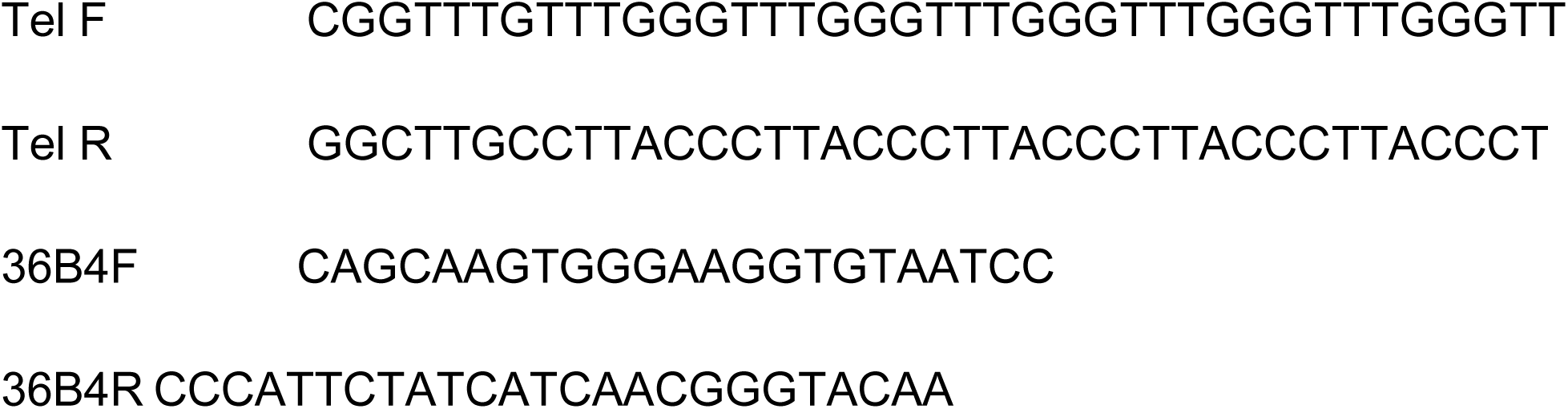

The relative telomere length of TNBC patient tissue samples were estimated by normalizing to telomere signal of HT1080 fibrosarcoma cells (reference cell type) as follows:

Telomere signal= 2 ^-(Ct Tel – Ct 36B4)^

Relative telomere length (RTL) = Telomere signal (sample) / Telomere signal (reference)

### Flow-cytometry

The procedure for Flow-cytometry was adapted from basic flow cytometry guidelines from CST and has been previously reported in Sharma *et al*,2021(Shalu Sharma et al., 2021). The cell suspension was fixed with 4% formaldehyde and permeabilized with 0.1% triton X (in 1XPBS) for nuclear proteins. Incubation of primary antibody (1:500 v/v in 1% BSA in 1X PBS) for 1 µg/ml antibody concentration for 2 hours at RT or at 4°Covernight, was followed by PBS wash (3 times) and fluorescently labelled secondary antibody ( 1:1000 v/v) for 2 hours at RT. Following this, cells were washed with 1X PBS (3 times) and re-suspended in 1XPBS for flow-cytometric analysis using BD Accuri C6 or BD FACSAria III. Equal number of events (10000 for most of the analysis) were recorded for calculating percentage infiltration.

For single channel analysis, histogram prolife (log scale) was generated in ungated sample-SSC-FSC plots. For quadrant gating for double positive cells (two channels), biex-scale plots or log scale were used. All statistics like mean fluorescence intensity and standard deviation were estimated using built-in tools for analysis in the software FlowJo.

### Relative Telomeres length (RTL) by Flow-cytometry

The RTL value is calculated as the ratio between the telomere signal of each sample and the control cell. The Telomere PNA Kit/FITC kit from Dako (Agilent) was used to estimate Telomere length signal in samples. The process has been previously standardized and reported (Mukherjee et al., 2018).

### TelSeq-estimation of telomere length using sequencing data

We used TelSeq(Ding et al., 2014) to estimate the average telomere length (TL) for each tumour and normal sample in our dataset. TelSeq was run with default parameters using WGS(whole genome Sequencing) samples with coverage ≥ 30X, except the read length parameter (-r) which was set to 150 to match with our data. Briefly, the TL was calculated as the total number of reads containing seven or more telomeric repeats (“TTAGGG”) divided by the total number of reads with GC content 48-52%. Further, the resultant value was multiplied with a constant value to account for genome size and the number of telomeric ends. This was done separately for each read group within a sample, and then the weighted average was calculated for each sample by taking into account the number of reads per group (as weights).

The estimated average TL [represented in Kilobase-pair (Kb) units] for tumour and normal samples has been provided in **Supplementary Figure 7B**. Overall, we found that the tumour samples have lower TL scores (Median: 3.20 Kb) as compared to the matched normal samples (Median: 4.67 Kb). Also, the TL estimates from our tumour samples (Median: 3.20 Kb, range:2.3 to 21.10 Kb) were comparable to the TCGA breast cancer cohort (WGS) (Median: 3.5 Kb, range: 1.4 to 29.7 Kb) analyzed by Barthel et al, 2017 (Barthel et al., 2017).

To check if the tumour samples with high TL estimates were confounded by the aneuploidy and tumour purity (which was not taken into account in the TelSeq analysis), we computed the purity and ploidy values for each tumour sample (where possible) using ASCAT(Raine et al., 2016) through Sarek pipeline(Garcia et al., 2020). However, we found no significant difference in the purity (Wilcoxon test two-sided *P =* 0.25) and ploidy (Wilcoxon test two-sided *P =* 0.52) values between tumour samples with high versus low TL scores, suggesting that the tumours with high TL estimates were not influenced by these parameters.

### IL1B and IL1RA treatment

Reconstituted IL1B (Sigma-Aldrich) was treated to appropriate cells at 10ng/ml concentration followed by assays. Similarly, IL1RA (Abcam) was treated to appropriate cell/TNBO at 20ng/ml concentration followed by assays at appropriate time points in specific experiments.

### IL1B ELISA

IL1B ELISA was performed using Human IL-1 beta/IL-1F2 DuoSet ELISA kit by R&D Biosystems. As per manufacturer’s protocol 96 well microplate was coated with100ul per well of freshly prepared Capture Antibody diluted 100 times in PBS without carrier protein. Thereafter the plate was sealed and incubated overnight at room temperature. The following day, the capture antibody was aspirated after which each well was washed three times with 1X Wash Buffer (PBST). After the last wash, the remaining wash buffer was removed carefully and completely by aspiration and blotting it against clean paper towels. The prepped, empty wells were blocked at room temperature for 1hr by adding 300 μL of Reagent Diluent (20% BSA in 1X PBST) to each well. Wash the wells as in the step above. Next we added the samples and standard protein of known increasing concentrations (100ul in volume) diluted in Reagent Diluent. The plates were then covered with an adhesive strip and incubated for 2 hours at room temperature. Post incubation, the wells were washed as above for plate preparation. Following the washes we added 100ul of freshly prepared Detection Antibody diluted 100 times in PBS. The plate was covered and incubated for 2hrs at room temperature. Post incubation the wells were washed thrice as in the previous step. Finally for generating detectable signal we added 100 μL of Streptavidin-HRP to each well, covered the plate and incubated it in dark at room temperature for 20mins. Next the wells were washed thrice as in steps above. After aspirating the wells clean of any buffer we added 100ul of Substrate Solution to each well and incubated the plates again in dark at room temperature for 20minutes. Finally, 50ul of stop solution was added to each well and mixed well by gentle tapping. The signal was measured using a microplate reader set to 450 nm, 570 and 540nm. Perform wavelength correction by subtracting readings at 540 nm or 570 nm from the readings at 450 nm.

### Antibodies

#### Primary antibodies

TRF2 rabbit polyclonal (Novus NB110-57130)- ChIP, IP, WB

TRF2 mouse monoclonal (Millipore 4A794)- Flow cytometry

TRF1 mouse monoclonal (Novus NB110-68281)-WB

IL1R1 rabbit polyclonal (abcam ab106278)- IF, Flow-Cytometry, WB

DDK/FLAG mouse monoclonal (Sigma-F1804)- ChIP, WB

BG4 recombinant antibody (Sigma Aldrich MABE917)- ChIP

P300 (CST D2X6N) rabbit monoclonal- ChIP, IF,WB

Ac-p300/CBP (CST 4771) rabbit monoclonal-ChIP,WB

CBP (CST 7389) rabbit monoclonal- ChIP,WB

P65(NFKB) (CST 8242) rabbit monoclonal- WB

Ser536 P p65 NFKB (CST 3033) rabbit monoclonal- WB

IKB (CST 4814) mouse monoclonal- WB

p-IKB (CST 2589) rabbit monoclonal- WB

CD44 (CST 3570) mouse monoclonal-IF

Histone H3 rabbit polyclonal (Abcam ab1791)-ChIP H3K27ac rabbit polyclonal (Abcam ab4729)-ChIP

H3K4me3 rabbit polyclonal (Abcam ab8580)- ChIP

H3K27me3 rabbit monoclonal (abcam ab192985)- ChIP

H3K9me3 rabbit polyclonal (abcam ab8898)- ChIP

Beta-actin mouse monoclonal (Santacruz C4)-WB

GAPDH mouse monoclonal (Santacruz 6C5)-WB

CD11b Monoclonal Antibody (ICRF44), eBioscience™- Flow cytometry

CD206 Monoclonal Antibody (MR5D3) Invitrogen- Flow cytometry

APC anti-human CD326 (EpCAM) (9C4) BioLegend- Flow cytometry

FITC anti-human CD11B (ICRF44) BioLegend- Flow cytometry

PE anti-human CD45 (HI30) BioLegend- Flow cytometry

PE-Cy7 anti-human CD45 (W17233E) BioLegend- Flow cytometry

Anti-Rabbit IgG/Anti-mouse IgG (Millipore) was used for isotype control.

#### Secondary antibodies

anti-Rabbit-HRP(CST), anti-Mouse-HRP(CST),mouse-ALP (sigma),rabbit-ALP(sigma) anti-rabbit Alexa Fluor 488, anti-mouse Alexa Fluor 594 (Molecular Probes, Life Technologies).

#### Real time PCR

Total RNA was isolated from cells using TRIzol Reagent (Invitrogen, Life Technologies) according to manufacturer’s instructions. For TNBC tissue and xenograft, Allprep DNA/RNA Universal kit from Qiagen was used. The cDNA was prepared using High-Capacity cDNA Reverse Transcription Kit (Applied Biosystems) using manufacturer provided protocol. A relative transcript expression level for genes was measured by quantitative real-time PCR using a SYBR Green (Takara) based method. Gene expression was calculated as 2^-delCT^ with delCT being the difference in threshold cycles (Ct) between test and internal control. *GAPDH* / 18S gene was used as internal control for normalizing the cDNA concentration of each sample. For fold change of gene expression calculations, 2^-delCT^ for the first replicate of the control sample was used for normalization for the samples. Normalized expression values (fold change) were plotted and compared with appropriate statistical testing.

RT PCR primers used have been enlisted as follows:

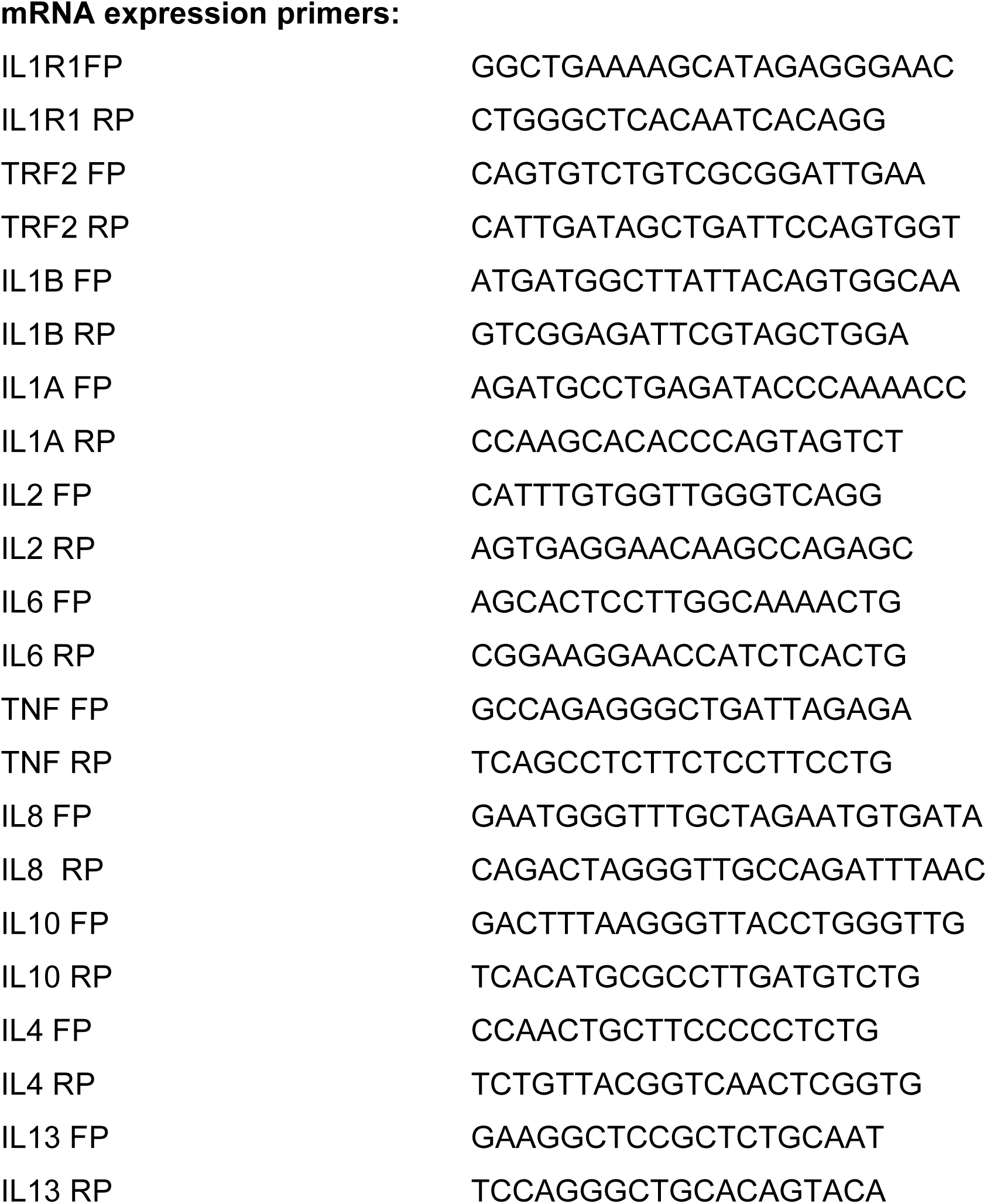
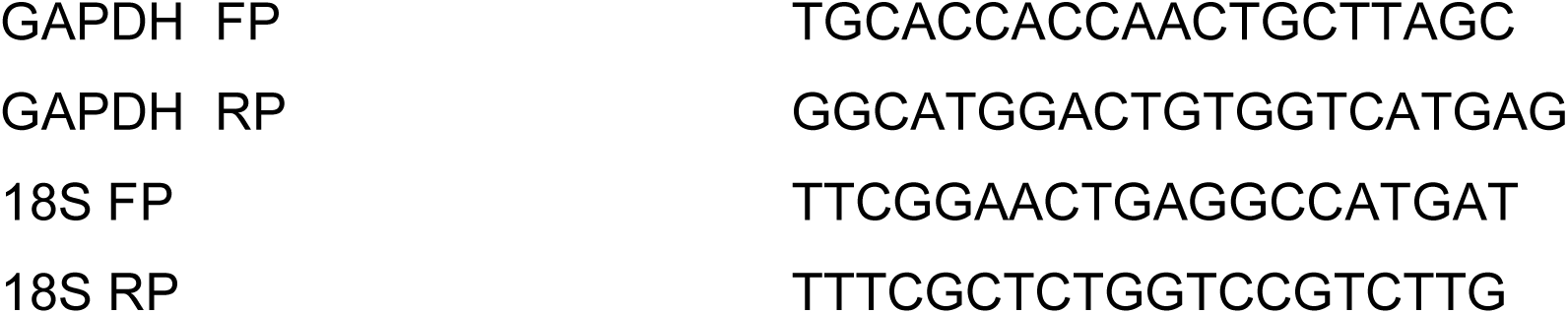

TRF2 target gene expression primers from Mukherjee *et al,* 2018 (Mukherjee et al., 2018).

#### RNA-seq analysis

RNA was extracted using TRIzol TruSeq RNA Library Prep Kit v2 was used from preparation of libraries for sequencing. Raw Illumina sequencing reads were checked for quality using FastQC (version 0.11.9) followed by adapter clipping and trimming using Trimmomatic (version 0.39)with default parameters. Trimmed reads were then aligned to the human reference genome (GRCh38, GENCODE v36) using STAR aligner (version 2.7.8a)(Dobin et al., 2013). FeatureCounts (subread package version 2.0.1) was used to quantify gene expression(Quinlan and Hall, 2010). Quality checks were performed at each step using the MultiQC tool (version 1.10.1). Differential gene expression analysis was performed using the DESseq2 package (version 1.30.0)(Love et al., 2016) in R (version 4.0.3). The analysis was performed by removing the effects of confounding variables such as batch and replicate using the appropriate design formula. Gene expression was normalized using variant stabilizing transformation (vst) for visualization purposes. Genes with BH-adjusted p value < 0.05 and absolute Log2 fold change greater than 1 (at least 100% fold change in either direction) were taken as significantly differentially expressed. Two replicates for each sample were used for differential gene expression analysis.

### ChIP (Chromatin Immunoprecipitation)

ChIP assays were performed as as previously reported by Mukherjee *et al*.,2018,2019 (Mukherjee et al., 2019a, 2018). Briefly, 3-5 million cells were fixed with ∼1% formaldehyde (Sigma-Aldrich) for 10 min and lysed. Chromatin was sheared to an average size of ∼300-400 bp using Biorupter (Diagenode). 10% of the sonicated fraction was processed as input using phenol–chloroform and ethanol precipitation. ChIP was performed for endogenous/ectopically expressed protein using 1:100 dilution (v/v) of the respective ChIP grade antibody incubated overnight at 4°C. Immune complexes were collected using Salmon sperm DNA-saturated Magnetic Dynabeads (-50 μg per sample) and washed extensively. Phenol-chloroform-isoamylalcohol was used to extract DNA from the immunoprecipitated fraction. ChIP DNA was quantified by Qubit HS ds DNA kit (Thermo Fisher Scientific). Quantified ChIP samples were validated by qRT-PCR.

ChIP enrichment was calculated over 1% input as : 2^-(CT^ ^ChIP-CT^ ^1%^ ^input)^. Following this, the ChIP enrichment was normalized to enrichment value for respective IgG (Mock) or total H3 ChIP.

The ChIP-qRT PCR primers are as follows:

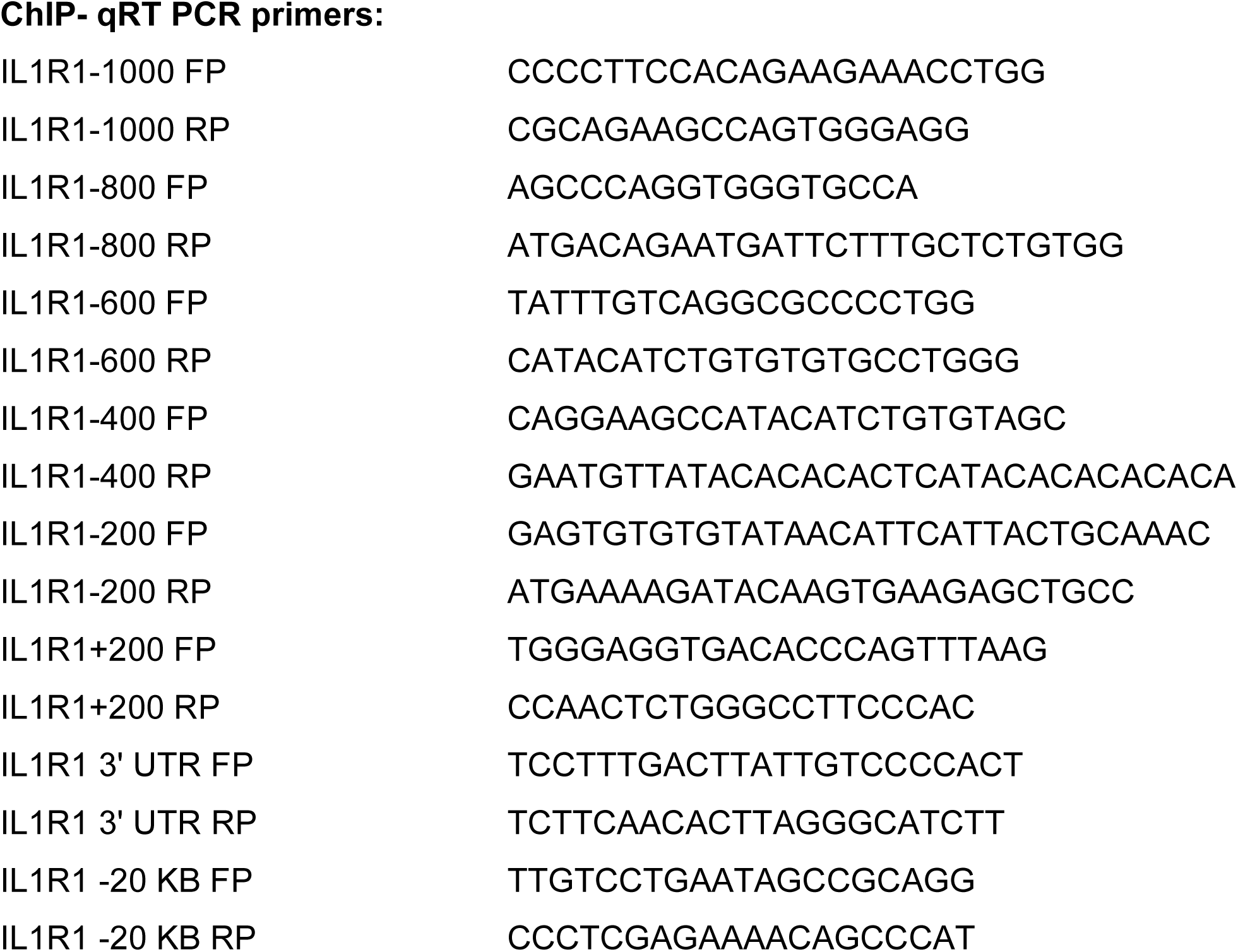

### Immunofluorescence microscopy

Adherent cells were seeded on coverslips and allowed to reach a confluency of ∼ 70%. Cells were fixed using freshly prepared 4% Paraformaldehyde by incubating for 10 min at RT. Cells were permeabilized with 0.25% Triton™ X-100 (10 min at RT) and treated with blocking solution (5% BSA in PBS) for 2 hrs at RT. All the above stated were followed by three washes with ice cold PBS for 5 mins each. Post-blocking, cells treated with relevant antibodies (1:500 for TRF2, 1:500 for IL1R1 and 1:1000 for CD44/p300) and incubated overnight at 4⁰C in a humid chamber. Post-incubation, cells were washed alternately with PBS and PBST three times and probed with secondary Ab (rabbit Alexa Fluor 488(1:1000) / mouse Alexa Fluor 594 (1:1000)) for 2 hrs at RT.

Cells were washed again alternately with PBS and PBST three times and mounted with Prolong Gold anti-fade reagent with DAPI. Images were taken on Leica TCS-SP8 confocal microscope/ Cytiva Deltavision microscope. Lasers of emission wavelength 405nm, 488nm and 532 nm were used imaging for DAPI, Alexa Fluor 488 secondary Ab and Alexa Fluor 594 Ab respectively. Laser intensity was kept 20% for DAPI and 40-60% for Alexa Fluor secondary antibodies. ImageJ (FIJI) software was used to calculate signal intensity (a.u.).

### Western blotting

For western blot analysis, protein lysates were prepared by suspending cell pellets in 1X RIPA (with 1x mammalian protease inhibitor cocktail). Protein was separated using 10%/12% SDS-PAGE and transferred to polyvinylidenedifluoride membranes (Immobilon FL, Millipore). After blocking, the membrane was incubated with primary antibodies. Post incubation with primary antibodies (1:1000 v/v), the membrane was washed with 1X PBS and then incubated with appropriate secondary antibodies (1:2000 v/v). Following secondary antibody incubation, the membrane was washed with 1XPBS. The blot was finally developed using HRP substrate/BCP NBT and reagents from Millipore.

### HT1080 TRF2 and TERT inducible cells

TRF2 cDNA sequence was inserted into pENTR11 vector and the cassette was transferred into lentiviral plasmid pCW57.1 by gateway cloning. Following this viral particles were generated from HEK293T cells and the viral soup with polybrene (Sigma-Aldrich) was used for transduction. Cells with insert were selected with 1μg/ml of puromycin and tested for TRF2 induction using varying doxycycline concentrations.

Similarly TERT cDNA was amplified from a mammalian expression vector and inserted into pENTR11 vector. Subsequently, the same strategy was used to generate and test TERT inducible HT1080 cells.

### Vector constructs

The pCMV6-myc-DDK(FLAG)-TRF2 vector (TRF2 WT) was procured from Origene (<u>RC223601</u>). The mutants delB and delM constructs used in the study have been previously reported(Hussain et al., 2017) and made through mutagenesis of the TRF2 WT construct. TRF2 shRNA was procured from Origene (<u>TL308880</u>). Side-directed Mutagenesis was performed on the TRF2 WT plasmid to generate the PTM mutants.

For customizing dCas9-TRF2 fusion plasmids, the backbone of dCAS9-VP64-GFP (Addgene 61422) was used. IL1R1 sgRNA was custom designed against IL1R1 promoter G-quadruplex (G4-motif B) and cloned in pX333 (Addgene 64073) backbone after replacing Cas9 with mCherry.

### Transfections

TRF2 wildtype (myc/DDK tag) or mutant mammalian expression vector pCMV6 was transfected into HT1080 /MDAMB231 cells that were 60 % confluent using Lipofectamine 2000 transfection reagent (following the manufacturers’ protocol). 2-4μg of plasmid was used for transfection in a 35 mm well for each case.

In case of TRF2 shRNA (Origene)/TERC shRNA (Santa Cruz Biotechnology), 4μg of plasmid was used for transfection in a 35 mm well for each case and Fugene HD transfection reagent was used as per manufacturer’s protocol. Cells were then maintained in 1μg/ml of puromycin for stable shRNA maintenance. For dCAS9-TRF2 fusion constructs, co-transfected with IL1R1 sgRNA plasmid, FugeneHD was used at recommended concentrations.

### TRF2 silencing by siRNA

HT1080 cells were transfected with TRF2 siRNA oligonucleotides (synthesized from Eurogenetics Pvt. Limited)(Shalu Sharma et al., 2021) using lipofectamine 2000 (Invitrogen) transfection reagent according to manufacturer’s instructions. Silencing was checked after 48 hr of transfection. Pooled scrambled siRNA was used as control.

### Luciferase assay

IL1R1 promoter (upto 1500 bp upstream of transcription start site) was cloned in PGL3 basic vector upstream of firefly luciferase construct. G4 motif disrupting mutations were introduced in the promoter construct by site directed mutagenesis.

Plasmid (pGL4.73) containing a CMV promoter driving Renilla luciferase was co-transfected as transfection control for normalization. After 48h, cells were harvested and luciferase activities of cell lysate were recorded by using a dual-luciferase reporter assay system (Promega).

### Circular Dichroism

Circular Dichroism profiles (220 nm to 300 nm) were obtained for representative G4 motifs identified within IL1R1 promoter and an overlap with TRF2 high confidence ChIP seqpeak. Circular Dichroism showed the formation of G4-motif with the unaltered sequence while mutated G4 sequence gave partial/complete disruption of the G4-motif under similar conditions (buffer used for G-quadruplex formation -10mM Na cacodylate and 100 mM KCl). The circular dichroism (CD) spectra were recorded on a Jasco-89spectropolarimeter equipped with a Peltier temperature controller. Experiments were carried out using a 1mm path-length cuvette over a wavelength range of 200-320 nm. Oligonucleotides were synthesized commercially from Sigma-Aldrich. 2.5uMoligonucleotides were diluted in sodium cacodylate buffer (10 mM sodium cacodylate and 100 mM KCl, pH 7.4) and denatured by heating to 95°C for 5 min and slowly cooled to 15°C for several hours. The CD spectra reported here are representations of three averaged scans taken at 20°C and are baseline corrected for signal contributions due to the buffer.

### CCR5-IL1R1 promoter insert cells

IL1R1 promoter and downstream firefly luciferase ( and G4 mutant variant ) was inserted in HEK293T cells by using strategy previously standardized and reported for TERT promoter-gaussia insertion in the same locus(Shalu Sharma et al., 2021).

### IL1R1 promoter G4 mutant HEK cells

Custom designed gRNA pair was used to insert mutant G4 template sequence (G4 mutantB) into the IL1R1 endogenous promoter by co-transfecting Cas9and gRNA pair with doner template SSODN sequence. Following this, single cell screen was done for positive mutant colony. Once identified an appropriate mutant colony was used for experiments.

sgRNA and SSODN details are as follows:

sgRNA1- 5’- GCTGCCAATGGGT**G**<u>GAGTC</u>TTG**G**G -3’ (efficiency score (Doench et al)- 35)

-ve strand, ssODN: 91bp (PAM proximal)- 36bp (PAM distal)

Target strand is +ve strand. Non-target stand is –ve strand. ssODN shud be +ve strand.

**ssODN** (mutated): chr2:102070276-102070402

GGGACACTGCAGCCCTGGCCTGGCCTACTTTTCTCTTCTCCCATATCTGGAAAATGAAAGT AGCCTGACGTATCCGGGGAC**A**CGGC**A**CAAGACTC**A**ACCCATTGGCAGCTCTTCACTTGTA TCTTTT

sgRNA2- 5’- GCAATGGGT**G**<u>GAGTC</u>TTG**G**GCCG**G** -3’ (efficiency score (Doench et al)- 40) -ve strand, ssODN: 85bp (PAM proximal)- 42bp (PAM distal)

### Immunoprecipitation

Immunoprecipitation of proteins was performed using modified version of suggested IP protocol from CST. Cells were lysed using cell lysis buffer ((1X) 20 mM Tris (pH 7.5), 150 mM NaCl (Sigma-Aldrich), 1 mM EDTA (Sigma-Aldrich), 1 mM EGTA (Sigma-Aldrich), 1% Triton X-100 (Sigma-Aldrich), 2X mPIC (Thermo Fisher Scientific)) and treated with relevant primary antibody (1: 100 v/v) overnight at 4°C or mock IgG on rotor. 50 µg of Dyanbeads (blocked with BSA-5% for 60 mins at RT) was added to the lysis soup and kept for 2 hrs in mild agitation. Dynabeads were magnetically separated and washed thrice (with 1X PBS with 0.1% Tween-20). The beads were resuspended with 1X RIPA (50 µl) and 1X Laemmli buffer. This was followed by denaturation at 95°Cfor 10 mins and followed by western blot for specific proteins.

### HAT assay

Histone acetyl transferase (HAT) assay was performed as per manufacturers’ protocol from Active motif. Mammalian TRF2 and p300 full length were commercially procured from origene and abcam respectively and used for the assay.

### IL1R1 knockout HT1080 cells

Two sets of gRNAs targeting either exon 3 or exon3-7 of *IL1R1* coding region were cloned in PX459 (Addgene 62988) and co-transfected in HT1080 cells, followed by puromycin selection and single cell screening for IL1R1 knockout cells. A knockout clone was selected (based on IL1R1 depletion) and used for further experiments.

### Gene ontology and cBIO PORTAL ANALYSIS

Gene Ontology for KEGG pathway enrichment and associated figures were generated using ShinyGO web portal(Ge et al., 2020). Gene ontology (Biological Process) analysis was done using gene list (Barthel et al., 2017) using publicly available resource –The Gene Ontology Resource database (http://geneontology.org/). Fold enrichment, FDR values and p-values were calculated using PANTHER. Ontology plot was made using MS Excel multivariate plot. *IL1R1* high/low samples were segregated, and differential gene expression gene list was generated using cBioPortal (https://www.cbioportal.org/) using the TCGA Breast Invasive carcinoma dataset (Firehose legacy). Sample IDs for the patient samples have been provided in **Supplementary file 4**. For *IL1R1* high samples, the cut-off used was 1.5 times the standard deviation of the mean expression value. Custom query commands were used to fetch the *IL1R1* high and low sample IDs. DEG list and survival plots were generated using in-built functions of cBioPortal.

### TRAP (Telomere Repeat Amplification protocol) for telomerase activity

Protein lysate was prepared in 1X CHAPS Lysis Buffer and the assay was performed using TRAPeze™ RT Telomerase Detection Kit (Merck) per manufacturer’s instructions.

### Statistical Analysis

All statistical tests were performed using interface provided within GRAPHPAD PRISM software. Median values and correlations were calculated using MS excel functions.

## Supporting information

revised manuscript source data and statistical tests

## Ethics statement

The authors declare that relevant ethical approvals were taken to procure and use human clinical patient tissue and mouse xenograft tissue for experiments whose results have been shared in the manuscript.

## Data availability

The authors declare that all data necessary to support the inferences are included in the manuscript and supplementary information. Additionally, raw data for the RNA sequencing (HT1080-HT1080-LT cells) has been made public (GSE267781).

## Acknowledgements

We acknowledge Manish Rai (confocal imaging), Debojyoti Chakraborty Lab (DeltaVision Microscopy), Vivek Rao (THP1 cells) and Mohit Agarwal (ELISA experiments) from CSIR-IGIB. Fellowships from CSIR (AKM, AS, SSR, SS, ASG, SVM, SB and DS); DBT-Wellcome Trust India Alliance (AKM, AS, MC, MV and AP); UGC (SD) and DST (DK) are acknowledged. R.S and N.G acknowledges intramural funding support from NCBS-TIFR. This work was supported by the DBT/Wellcome Trust India Alliance Fellowship [grant number IA/S/18/2/504021] awarded to SC, and research funding from the Council of Scientific and Industrial Research (CSIR), Ashoka University and Department of Biotechnology (DBT).

## Author Contributions

Experimental design and execution: Tel-PNA FISH –SD; Tel-qRT PCR- MC,MV and AKM; Tel- Flow- AKM;CD206 IHC-AS and AKM;CD206/CD11b Flow- AKM and MC;TNBO-M2 infiltration flow-AKM; gene expression(TNBC)- AKM and MC, Telomerase activity-MC, gene expression(TNBO, xenograft, ex-vivo)-AKM,SS. IF-AKM,SVM; ChIP-AKM,SS,ASG,AS; Luciferase assays- AKM,SSR,ASG; WB- AKM,SVM,ASG,SSR,SD;IP-AKM;HAT Assay- AKM; IL1B ELISA- SS,AKM; gene expression(dCAS9 related)-SD; Immuno-Flow for IL1R1- AKM,SD; Immuno-Flow NFKB related- AKM,SD; Data Analysis and pipeline development- AB,SKB,IG,NG,RS,SD,AKM;

Resources: TNBC tissue collection- AS; TNBO generation- DK, AKM,SVM,AS,AP; THP1- Macrophage–AKM; NOD-SCID xenograft tumours-AKM,SD; Luciferase constructs-SSR, ASG; CCR5-insert cells-SSR,AKM; HEK IL1R1 G4 mutant cells- SSR,AS; IL1R1 gRNA- SSR; HT1080- TERT inducible- ASG,DS;HT1080-TRF2 inducible- ASG; HT1080 IL1R1 KO- SSR,AKM; HT1080 IL1R1 KO TRF2 inducible- AKM;MDAMB-231 LT cells-SS,AP; TRF2 PTM mutant plasmids- SB; TRF2 protein extraction-SB and ASG; TNBC whole genome sequencing-AS;HT1080-HT1080-LT RNA seq- SS, VA, MF;

Data collation-AKM, SD, AS, SSR, ASG, SVM, SC; Figure preparation-AKM, SD, AS, SC; Manuscript writing and editing- AKM, SD, AS, SC; Conceptualization of idea and planning of experiments- AKM and SC; Overall supervision and funding-SC.

## Inclusion and diversity statement

*We support inclusive, diverse, and equitable conduct of research*.

## Materials & Correspondence

*All correspondences should be addressed to SC*

---

**Additional supplementary figure 1.**
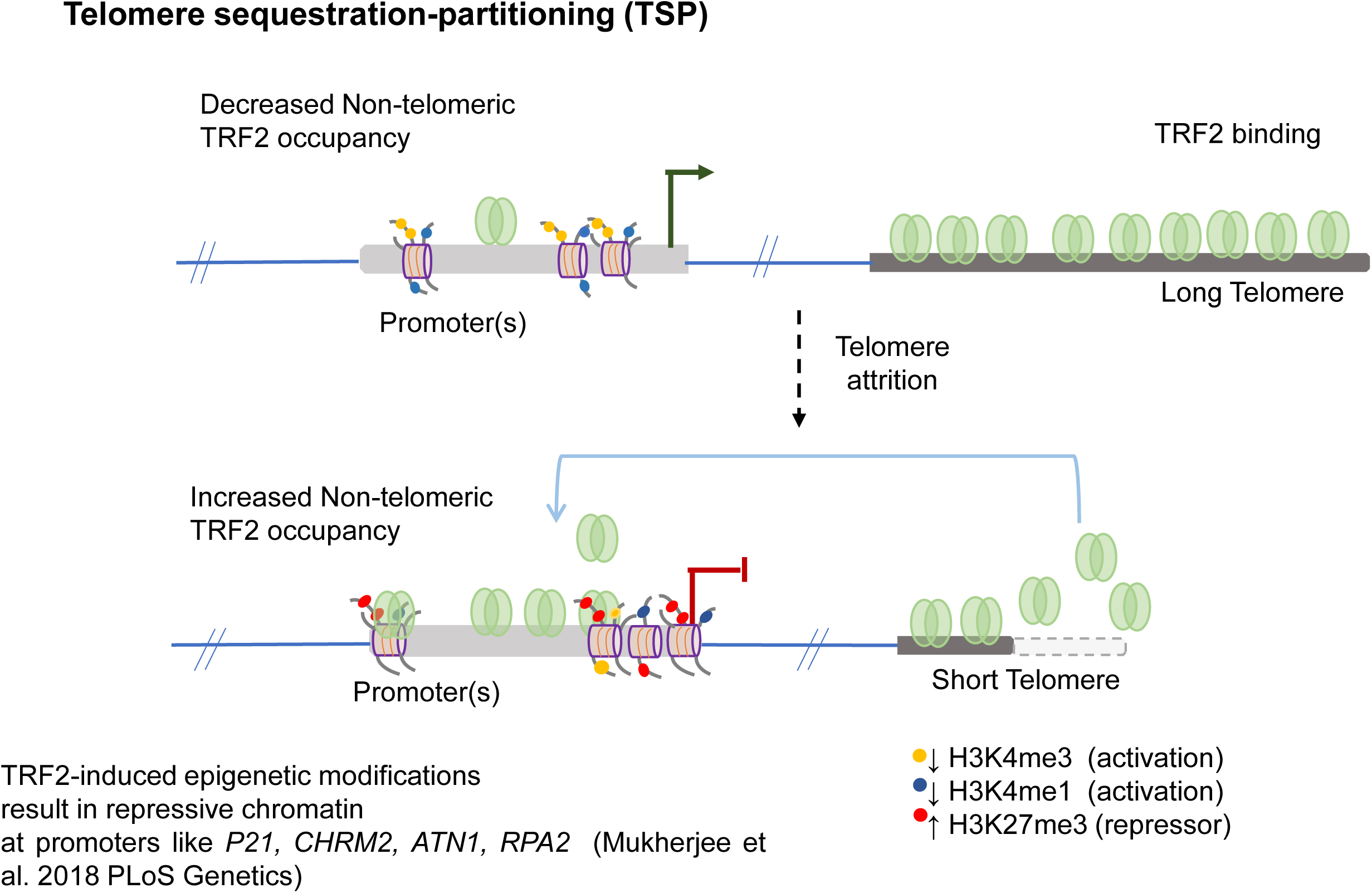
Graphical representation of the Telomere Sequestration Partitioning (TSP) model. The binding of TRF2 outside of telomeric regions (non-telomeric) is dependent on telomere length. When telomeres shorten, the presence of TRF2 at telomeres diminishes, resulting in increased TRF2 binding at non-telomeric sites. This shift in TRF2 distribution triggers epigenetic modifications at promoters showing telomere-dependent gene regulation. Described in Mukherjee et al. *PLoS Genetics*, 2018.

**Supplementary Figure 1:**
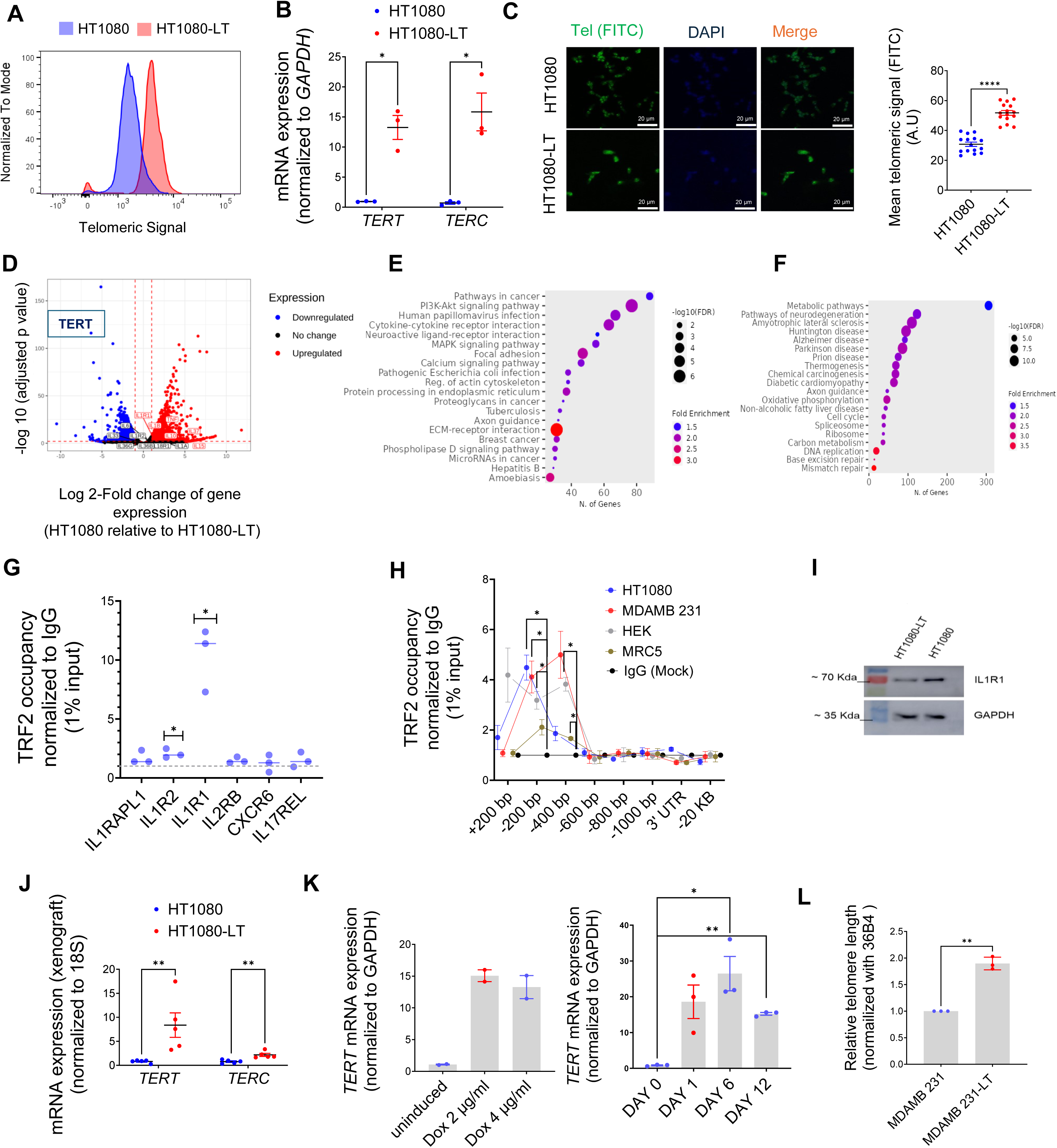
A. Telomer length (TL) by flow-cytometry analysis of telomeric signal (FITC labeled PNA probe) showing increase in TL in HT1080-LT cells relative to HT1080 cells; plotted along x-axis in log scale. B. *hTERT* and *TERC* expression as expected was enhanced in HT1080-LT relative to HT1080 cells (normalized to GAPDH). [N=3] Statistical significance was calculated using unpaired T test with Welch’s correction (p values: * ≤ 0.05, ** ≤ 0.01, *** ≤ 0.001, **** ≤ 0.0001). C. Immuno-histochemical (IHC) staining of previously reported and well-characterized cells with short/long telomeres (fibrosarcoma HT1080 or the long-telomere version HT1080-LT cells in cell-isogenic background) hybridized with telomere-specific FITC-labelled PNA probe, counterstained with DAPI ( left panel); telomeric signal was enhanced in HT1080-LT cells (quantified in right panel). [N=15] Statistical significance was calculated using Mann-Whitney’s non-parametric test (p values: * ≤ 0.05, ** ≤ 0.01, *** ≤ 0.001, **** ≤ 0.0001). D. Volcano plot of expressed genes in RNA-seq in HT1080 or HT1080-LT cells; x-axis has log-2 fold change for genes and the y-axis represents adjusted p values in log-10 scale. Key inflammation-related genes that have been marked in figure – most differentially expressed genes like *IL1R1*,*IL1B* and *IL10* were found to have higher expression in HT1080 cells compared to HT1080-LT cells. TERT which ( as expected) is significantly lower in HT1080 cells compared to HT1080-LT cells have been also marked. Upregulated and downregulated genes listed in supplementary file 1. E. Gene Ontology (KEGG) for upregulated genes in HT1080 cells over HT1080-LT cells. F. Gene Ontology (KEGG) for down-regulated genes in HT1080 cells over HT1080-LT cells. G. Occupancy of TRF2 by ChIP-qPCR checked in HT1080 cells for cytokine-related genes. Enrichment was plotted relative to mock IgG (post normalization to 1% input ) [N=3] Statistical significance was calculated using unpaired T test with Welch’s correction (p values: * ≤ 0.05, ** ≤ 0.01, *** ≤ 0.001, **** ≤ 0.0001). H. TRF2 occupancy on the *IL1R1* promoter in HT1080, MDAMB231, HEK293T or MRC5 cells by chromatin-immunoprecipitation (ChIP) followed by qPCR. Primers spanning +200 to -1000 bp of TSS used for TRF2 enrichment on the gene promoter; 3’UTR and -20 Kb upstream of TSS used as negative control for TRF2 binding; fold-change of occupancy was calculated over mock IgG after normalizing signal to 1% input. [N=3] Statistical significance was calculated using unpaired T test with Welch’s correction (p values: * ≤ 0.05, ** ≤ 0.01, *** ≤ 0.001, **** ≤ 0.0001). I. IL1R1 protein levels in HT1080 and HT1080-LT cells by western blot ; GAPDH was used as loading control. J. mRNA expression of *hTERT* and *TERC* (normalized to 18S) in xenograft tumors in NOD-SCID mice injected with HT1080 or HT1080-LT cells. *18S* has been used as a normalizing control. [N=5] Statistical significance was calculated using Mann-Whitney’s non-parametric test (p values: * ≤ 0.05, ** ≤ 0.01, *** ≤ 0.001, **** ≤ 0.0001). K. *hTERT* mRNA expression in hTERT-inducible HT1080 cells on treatment with different concentrations of doxycycline [N=2] (left panel) and at different days following induction [N=3] (right panel). *GAPDH* was used for normalization. Statistical significance was calculated using unpaired T test with Welch’s correction (p values: * ≤ 0.05, ** ≤ 0.01, *** ≤ 0.001, **** ≤ 0.0001). L. Relative telomere length by qRT-PCR in MDAMB231 or MDAMD231-LT (long telomere) cells in three independent experiments. [N=3] Statistical significance was calculated using unpaired T test with Welch’s correction (p values: * ≤ 0.05, ** ≤ 0.01, *** ≤ 0.001, **** ≤ 0.0001). Error bars correspond to standard error of mean from independent experiments.

**Supplementary Figure 2:**
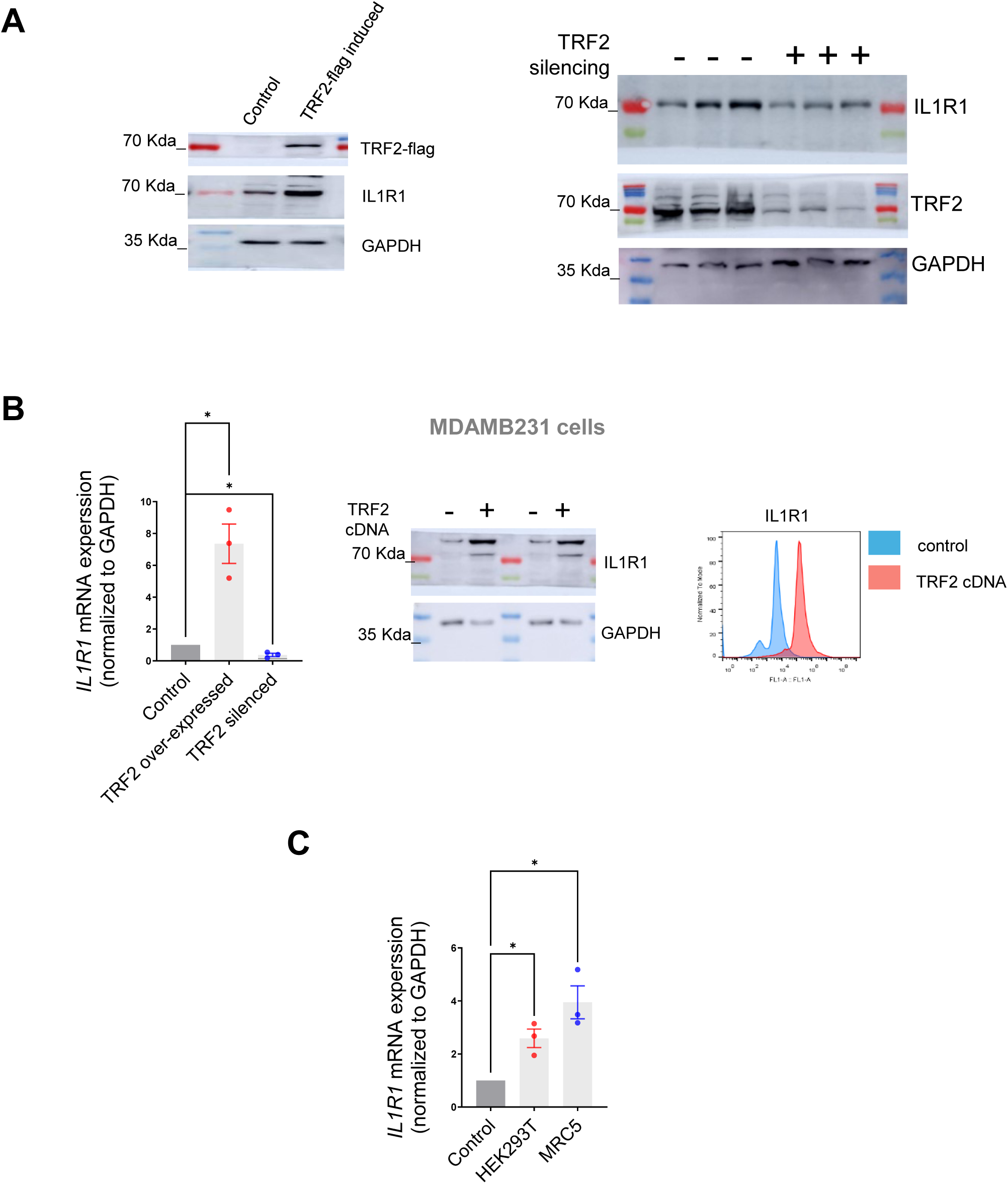
A. IL1R1 protein levels on TRF2-flag induction (left) or TRF2 silencing (right) in HT1080 cells following 48 hrs. of transfection. B. IL1R1 mRNA expression in MDAMB231 cells following expression of TRF2 cDNA or TRF2 silencing for 48 hrs; GAPDH used as normalizing control (left). [N=3] Statistical significance was calculated using unpaired T test with Welch’s correction (p values: * ≤ 0.05, ** ≤ 0.01, *** ≤ 0.001, **** ≤ 0.0001). IL1R1 protein levels in MDAMB231 cells following TRF2 induction for 48 hrs by western blot (centre) and immuno-flow cytometry (right). C. *IL1R1* mRNA expression in HEK293T and MRC5 cells following transfection of TRF2-cDNA for 48 hrs; *GAPDH* was used as normalizing control. [N=3] Statistical significance was calculated using unpaired T test with Welch’s correction (p values: * ≤ 0.05, ** ≤ 0.01, *** ≤ 0.001, **** ≤ 0.0001). Error bars correspond to standard error of mean from independent experiments.

**Supplementary Figure 3:**
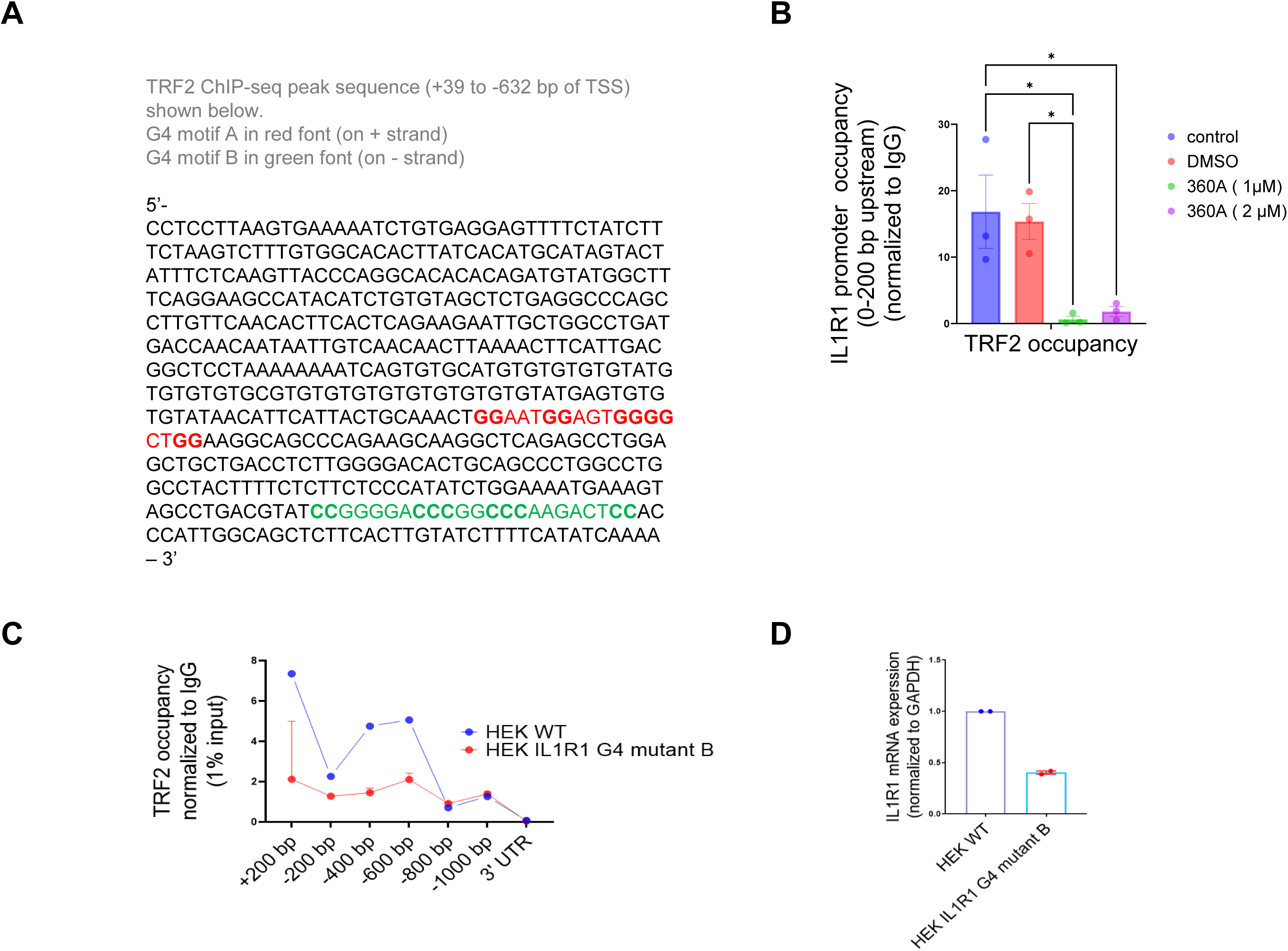
A.TRF2 ChIP-seq peak with sequence of the G4 motifs (A and B) and their respective positions on the *IL1R1* promoter B. Occupancy of TRF2 by ChIP-qPCR at the *IL1R1* promoter at the region of 0-200 bp of TSS in presence of ligand 360A in three independent experiments in HT1080 cells. The dosage of 360A used has been previous standardized (Mukherjee et al, 2019) [N=3] Statistical significance was calculated using Tukey’s multiple comparisons test (p values: * ≤ 0.05, ** ≤ 0.01, *** ≤ 0.001, **** ≤ 0.0001). C-D. Base substitutions for the G4 motif B in the IL1R1 promoter (as shown in Figure 3D) were introduced using CRSIPR in HEK293T cells. TRF2 ChIP-qPCR spanning the IL1R1 promoter (left) and *IL1R1* mRNA expression (right) in HEK293T cells with or without the G4-disrupting substitution in two replicates. [N=2] Error bars correspond to standard error of mean from independent experiments.

**Supplementary Figure 4:**
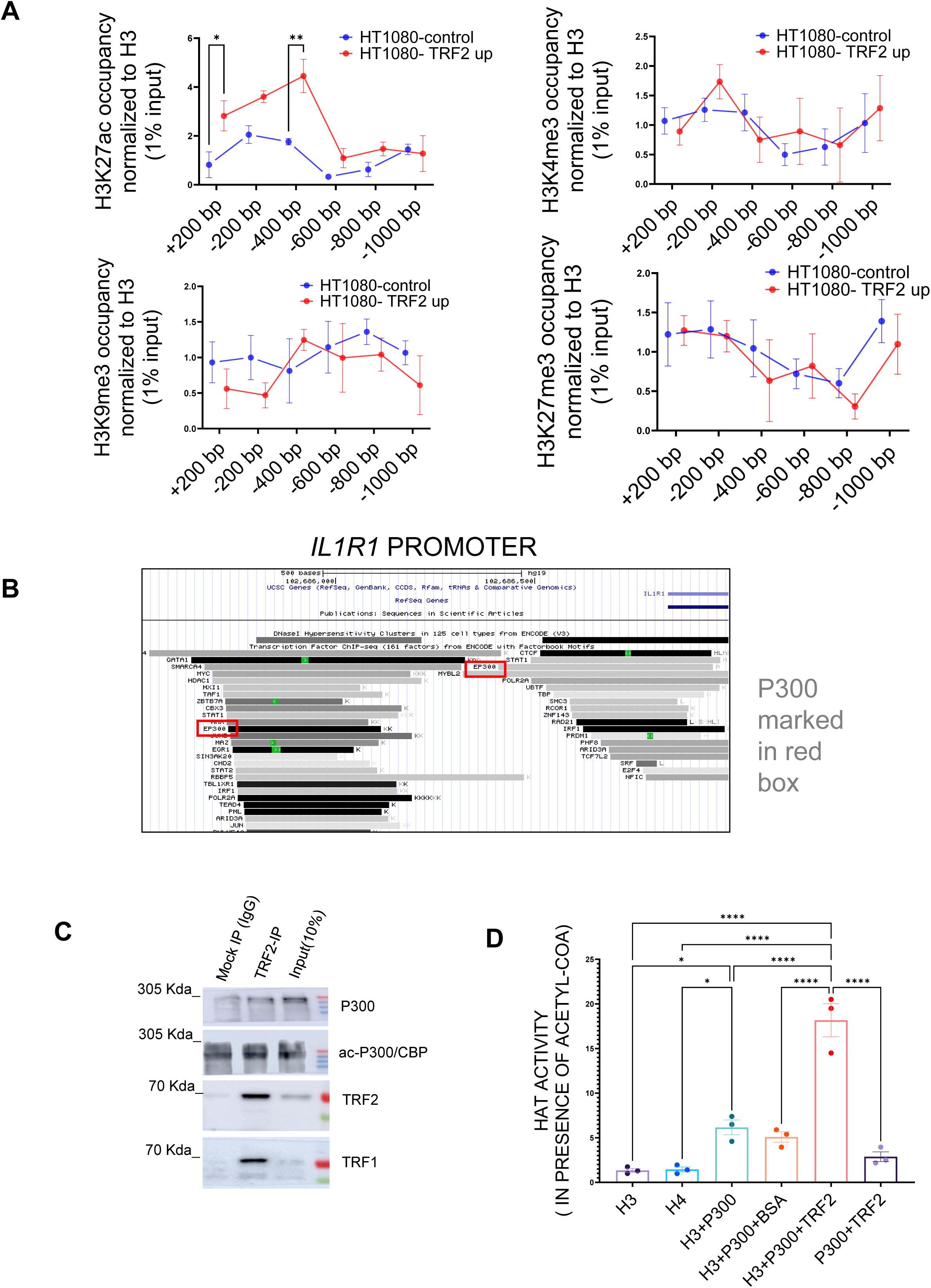
A. Enrichment of histone marks H3K27ac, H3K4me3, H3K27me3, H3K9me3 (normalized to total H3) by ChIP-qPCR on the *IL1R1* promoter in HT1080 cells in TRF2-up (induced) condition or un-induced corresponding control cells. [N=3] Statistical significance was calculated using Šídák’s multiple comparisons test (p values: * ≤ 0.05, ** ≤ 0.01, *** ≤ 0.001, **** ≤ 0.0001). B. Transcription factors/histone remodelers on the IL1R1 proximal promoter (-750 bp) across seven cell lines curated from ENCODE was plotted using UCSC genome browser hg19 human genome assembly. P300 enrichment marked in red box. C. Immunoprecipitation using TRF2 antibody probed for p300, acp300/CBP, TRF2 or TRF1 (positive control) in input, TRF2 IP fraction or mock IP fraction in HT1080 cells. D. Histone acetyl transferase (HAT) assay using purified histone H3/H4 and full-length p300 as substrate in presence/absence of recombinant TRF2; BSA (bovine serum albumin) was used as a non-specific mock protein. [N=3] Statistical significance was calculated using Tukey’s multiple comparisons test (p values: * ≤ 0.05, ** ≤ 0.01, *** ≤ 0.001, **** ≤ 0.0001). Error bars correspond to standard error of mean from independent experiments.

**Supplementary Figure 5:**
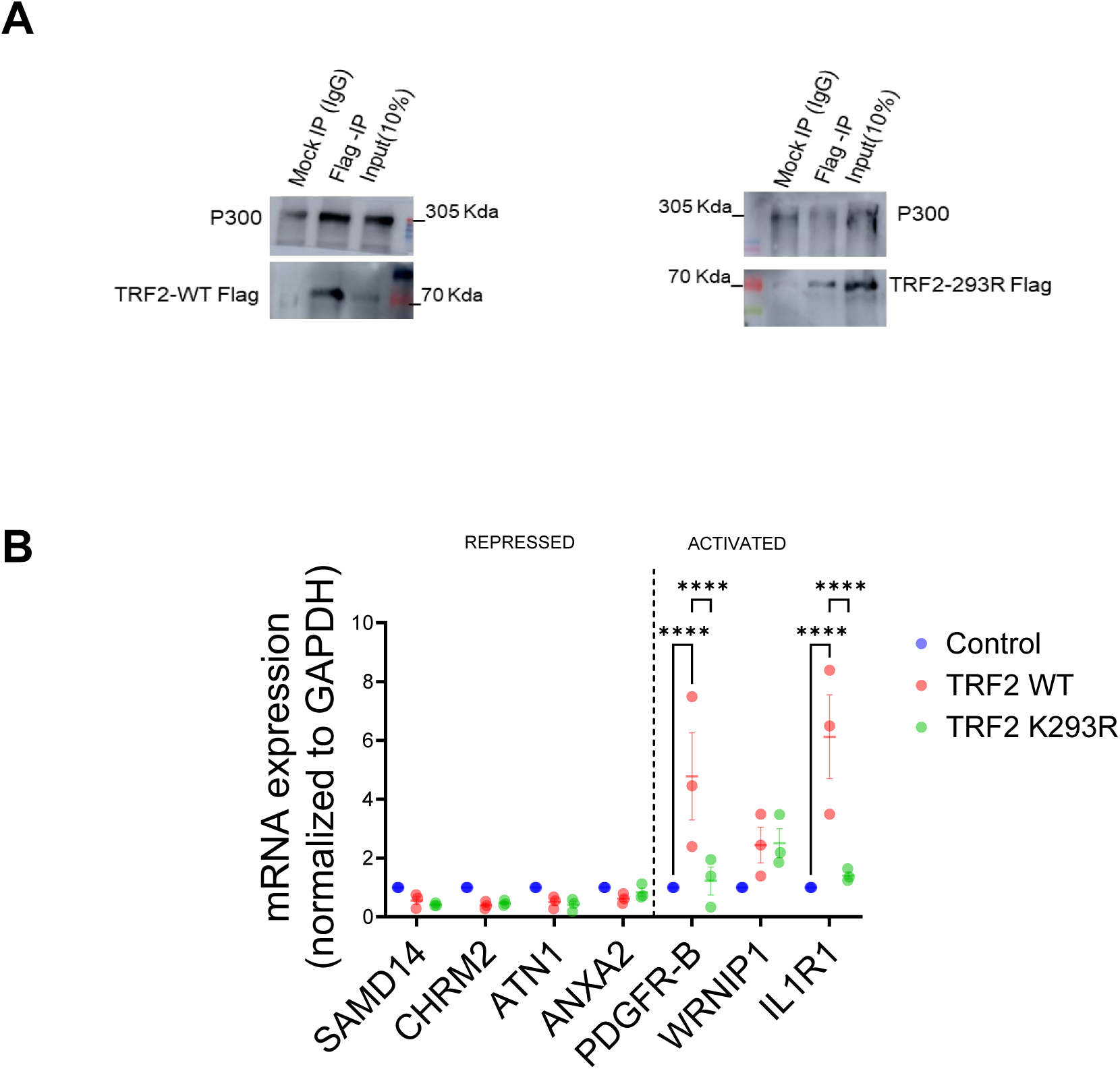
A. TRF2-WT Flag or TRF2 K293R-Flag was stably over expressed in HT1080 cells and immunoprecipitation was performed using Anti-Flag antibody or Mock IgG. Enrichment for p300 in IP faction was tested for TRF2-WT (left) and TRF2 K293R(right) B. Previously reported TRF2 target genes (Mukherjee et al, 2018) were checked following transient over-expression of TRF2 WT and TRF2 K293R in HT1080 cells. While K293R did not affect TRF2 mediated repression, it affected TRF2 activated genes PDGFR-B and IL1R1 but not WRNIP1. [N=3] Statistical significance was calculated using Tukey’s multiple comparisons test (p values: * ≤ 0.05, ** ≤ 0.01, *** ≤ 0.001, **** ≤ 0.0001). Error bars correspond to standard error of mean from independent experiments.

**Supplementary Figure 6:**
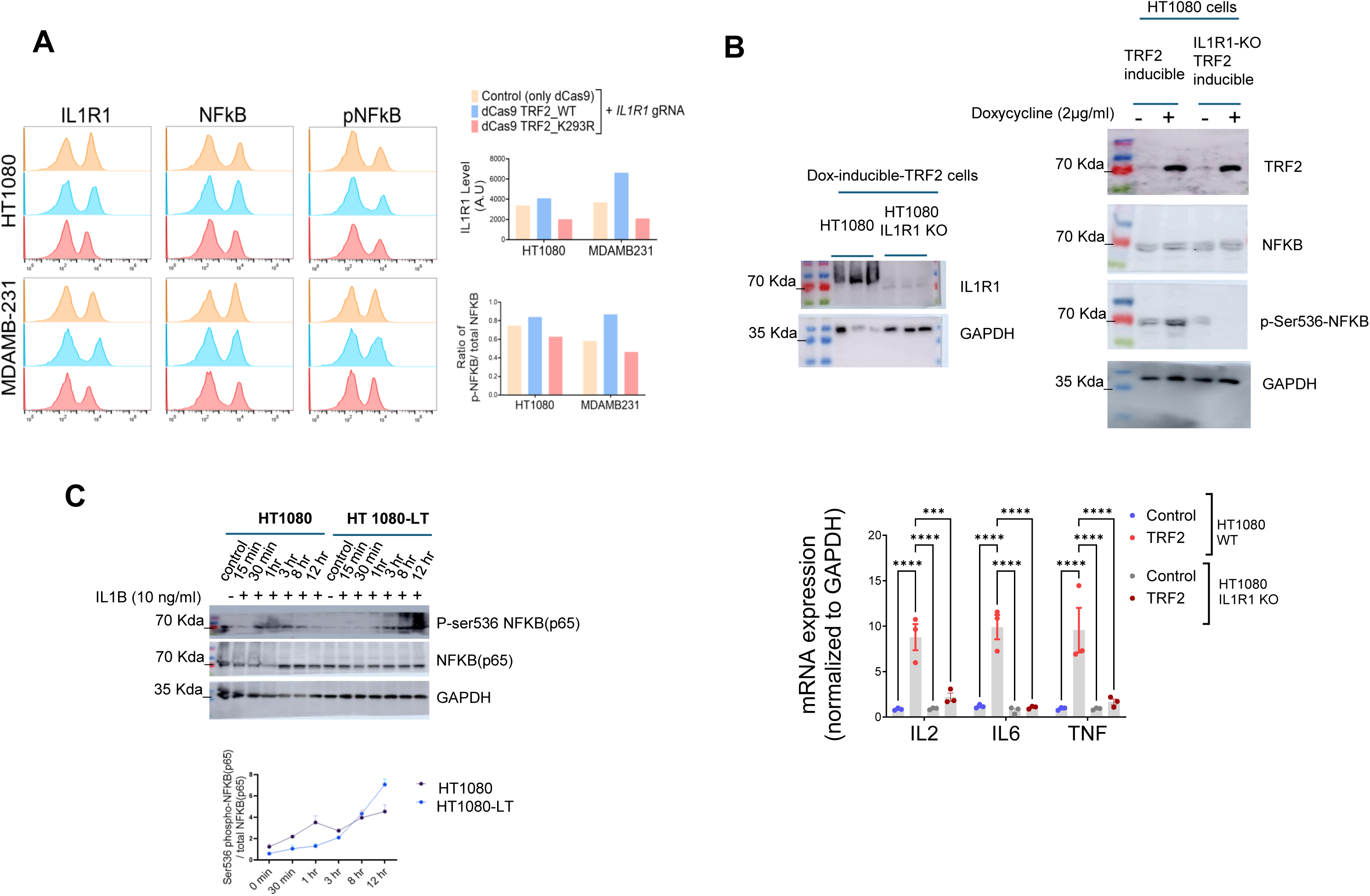
A. TRF2-wild type (WT) and TRF2-2293R mutant proteins fused with dCAS9 expressed and targeted to the IL1R1 promoter using IL1R1-specific gRNA in HT1080 (left) or MDAMB231 cells (right). Following this, NFKB and p-NFKB (Ser536-phosphorylation) levels were assessed by flow-cytometry in ten thousand HT1080 or MDAM231 cells ; mean fluorescence signal for IL1R1 and ratio of p-NFKB/total NFKB plotted in graphs on right panels. [N=1] B. IL1R1 knockout (KO) HT1080 cells using CRISPR were made and were transduced (lentiviral) to a doxycycline-inducible-TRF2 stable line (top left). TRF2, NFKappa-B and Ser536p-NFKappa-B in IL1R1 knockout or control HT1080 cells with or without TRF2 induction by doxycycline; GAPDH used as loading control (top right). Induction of TRF2 in HT1080 WT and IL1R1 KO cells to check for NFkappaB downstream target gene expression(bottom). Activation of NFKB target genes (∼ 2-5 fold) post-TRF2 over-expression was lower in IL1R1KO cells compared to WT cells (∼ 10 fold ) [N=3] Statistical significance was calculated using Tukey’s multiple comparisons test (p values: * ≤ 0.05, ** ≤ 0.01, *** ≤ 0.001, **** ≤ 0.0001). C. Phospho-NF-Kappa-B-(Ser-536p) and NFkappaB in HT1080 or HT1080-LT cells with or without IL1B stimulation (top panel); ratio of p-NFKB to total NFKB from above western blots plotted from two independent experiments (bottom panel). The NFkappa B (p65/RELA) is detected just below the 70 Kda molecular weight marker. Error bars correspond to standard error of mean from independent experiments.

**Supplementary Figure 7:**
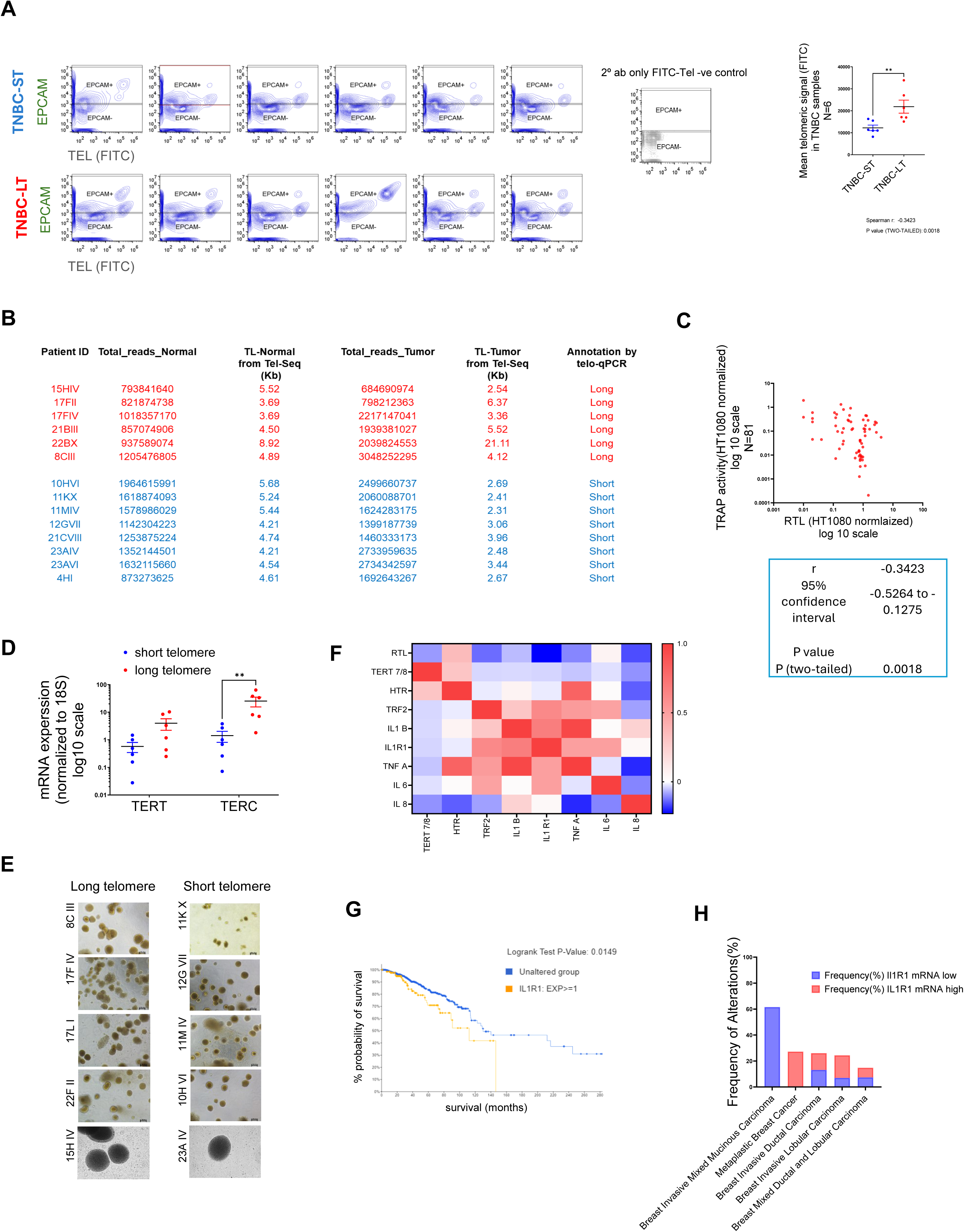
A. TNBC samples (N= 6 each for long or short telomere samples) stained with EpCAM (far red) and then hybridized with telomere specific PNA probe (FITC). Mean telomeric signal was plotted in total cells. [N=6] Statistical significance was calculated using Mann-Whitney’s non-parametric test (p values: * ≤ 0.05, ** ≤ 0.01, *** ≤ 0.001, **** ≤ 0.0001). B. Telomere length was determined using the previously published algorithm Tel-Seq from sequenced genomes of TNBC samples (N=8 for TNBC-ST (short telomere) and N=6 for TNBC-LT (long telomere); and adjacent normal tissue from same patient). Table with sequencing related details of the normal/tumor samples has been provided. C. Telomerase activity was measured from protein lysates across 64 TNBC samples (long or short telomeres) normalized to the telomerase activity of HT1080 cells for the same amount of protein; spearman correlation was calculated between RTL (relative telomere length) and telomerase activity. D. mRNA expression of *hTERT* and *TERC* in six TNBC samples (long or short telomeres) normalized to the *18S* gene in respective samples. [N=6] Statistical significance was calculated using Mann-Whitney’s non-parametric test (p values: * ≤ 0.05, ** ≤ 0.01, *** ≤ 0.001, **** ≤ 0.0001). E. TNBC organoids (TNBO) formed from TNBO-ST (short telomere) and TNBO-LT (long telomere) samples. F. Spearman correlation of,*TERT*, *TERC, IL1R1* and cytokine related gene expression and RTL (Relative telomere length) in 34 TNBC tissue samples with each other and RTL represented as heat map; gene expression was normalized to the18S gene; for telomere length the 36B4 single-copy gene was used for normalization (as in Figure 1D); fold change of TL was calculated with respect to TL in HT1080 cells. G. Overall Survival for breast cancer patients (BRCA) with high IL1R1 mRNA expression [N=254] or low IL1R1 mRNA expression [N=709] from TCGA datasets ( see Supplementary file 4 for details ) – graph tabulated using cBIOPORTAL with custom queries. Logrank p-value was found to be 0.0149. H. Frequency of occurrence of samples with high / low IL1R1 mRNA expression in various annotated sub-types of breast cancer from TCGA ( see Supplementary file 4 for details ) Error bars correspond to standard error of mean from independent experiments.

**Supplementary Figure 8:**
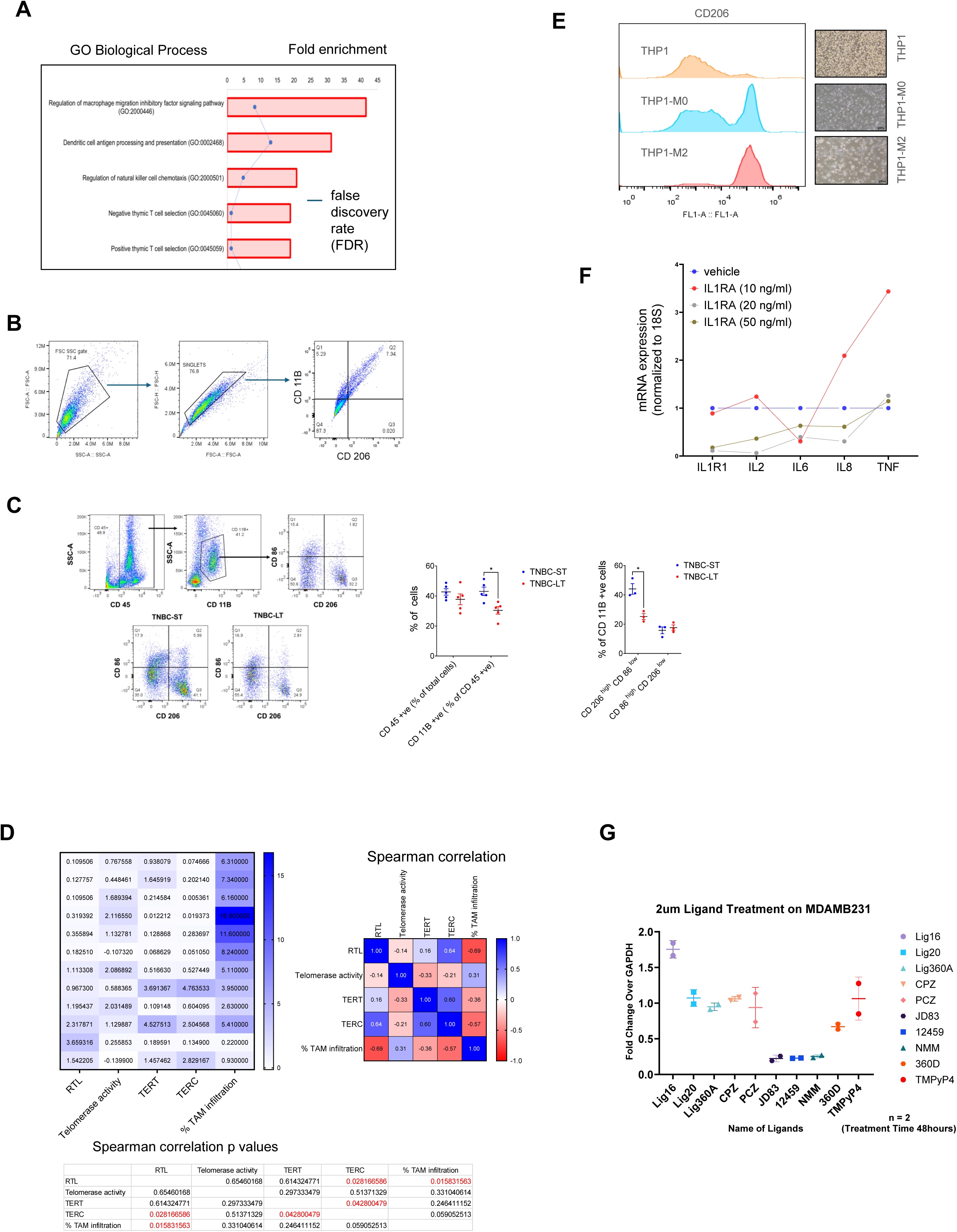
A. Gene Ontology (GO) analysis of top 500 genes correlated with telomere elongation across 31 cancer types( as reported in Barthel et al., 2017) Top 5 hits for ‘biological processes’ represented in descending order of enrichment; respective fold enrichments shown by individual bars (x-axis – top); the false discovery rate (FDR) shown by blue line. B. Gating strategy for Figure 8B. C. Gating strategy for CD 45 +ve c gated ells and CD 11B +ve gated cells in TNBC. Percentage of CD 45 +ve and CD11B +ve ( in CD 45 +ve population ) was checked for TNBC-ST and TNBC-LT samples. [N=5] Statistical significance was calculated using Mann-Whitney’s non-parametric test (p values: * ≤ 0.05, ** ≤ 0.01, *** ≤ 0.001, **** ≤ 0.0001).. In the CD11B +ve ( in CD 45 +ve population ), CD 206 (M2 macrophage marker) and CD 86 (M1 macrophage marker ) +ve cells were quantified. Percentage of CD 206 high CD 86 low ( M2 macrophages) and CD86 high Cd206 low (M1 macrophages ) in the myeloid cell population ( CD 45 +ve CD 11B +ve ) was plotted for three TNBC-ST and TNBC-LT samples each. [N=3] Statistical significance was calculated using unpaired T test with Welch’s correction (p values: * ≤ 0.05, ** ≤ 0.01, *** ≤ 0.001, **** ≤ 0.0001). D. TNBC tissue (N=6 each for long or short telomere samples) were tested for TERT and TERC expression (The Telomerase activity for these samples was estimated with HT1080 cell lysate as a reference. The normalized values ( to the group average) were plotted in a heat map (left) for telomerase activity, TERT and TERC gene expression in the TNBC samples used in Supplementary Figure 7D along with corresponding % TAM values (top left). The Spearman correlation matrix (top right) and p values (bottom).have been provided. E. CD206 levels by immuno-flow cytometry in THP1 cells and macrophages differentiated from THP1 cells into M0 or M2 types (described in Methods) and plotted along X-axis in log scale (top panel). Bright-field images at 10X magnification for each cell stage shown in top right panel. F. Different dosages of IL1RA was checked based on previously published reports and the minimal concentration where change in IL1R1 expression was seen ( 20 ng/ml) was used in other functional assays. [N=1] G. mRNA expression of IL1R1 was tested in MDAMB-231 cells after a 48-hr. treatment with various reported G4-binding ligands. *GAPDH* was used as a normalizing control. [N=2] Error bars correspond to standard error of mean from independent experiments.

